# Cell-of-origin shapes tumor subtype in *BRCA1*-associated triple-negative breast cancer

**DOI:** 10.64898/2026.09.29.755499

**Authors:** Tatyana Lev, Angela Lincy Prem Antony Samy, Sharmila Mallya, Isam Adam, Dana Pueschl, Dennis Ma, Timothy P. McMullen, Mandeep S. Lev, Jessica G. Gonzalez, Aaron Longworth, Arjun Nair, Xinyi Lu, Andrew K. Godwin, Paul A. James, Phuong L. Mai, Kenneth Offit, Heather Thorne, Antonis C. Antoniou, Georgia Chenevix-Trench, Susan M. Domchek, X. Steven Chen, Arthur D. Lander, John Lowengrub, Katherine L. Nathanson, Kai Kessenbrock, Devon A. Lawson

## Abstract

Pathogenic germline *BRCA1* variants greatly increase lifetime breast cancer risk, with no targeted prevention. Accrual of mutations following DNA repair errors in luminal progenitors is considered the chief mechanism of triple-negative breast cancer (TNBC) initiation, which cannot explain the organ specificity nor the phenotypic spectrum of the disease. Here we show that TNBC also originates in basal-myoepithelial cells, and that the cell of origin shapes TNBC subtype through aberrant epithelial differentiation, likely due to the pleiotropic role of BRCA1 in regulating hormone signaling. Single-cell RNA sequencing of 67,001 cells from *Wap-Cre Brca1/p53* mice and reanalysis of published *Blg-Cre* data show that aberrant basal-myoepithelial differentiation (*Wap-Cre*) generates the mesenchymal subtype (TNBC-M) and aberrant luminal differentiation (*Blg-Cre*) the basal-like subtype (TNBC-BL1). We also show that both subtypes occur at comparable frequency in *BRCA1* carriers. Aberrant basal-myoepithelial differentiation precedes tumor formation, and only tumors derived from transplanted basal cells reproduce the basal-containing phenotype of spontaneous tumors. Comparison with a developmental atlas identifies de-differentiation of myoepithelial cells into basal cells lacking myoepithelial markers, a fetal state, whereas experimental metastases shift toward luminal states while retaining a TNBC-M-like transcriptional program. These findings support the inclusion of basal-myoepithelial and luminal cells in targeted prevention strategies for *BRCA1* carriers, and adoption of TNBC subtyping for patient stratification and improving outcomes.

## Introduction

Germline pathogenic variants in *BRCA1* strongly predispose to breast cancer, with a cumulative risk of up to 72% by age 80^1^. BRCA1 functions in cell-cycle checkpoint control and homologous recombination, and its loss forces cells into error-prone repair pathways, generating the large-scale copy-number gains and losses that characterize *BRCA1*-mutant breast cancers^2–8^. Genomic instability alone does not account for the tissue restriction of the phenotype, as carriers develop cancer predominantly in hormone-responsive organs. BRCA1 also targets hormone receptors for degradation through its E3 ubiquitin ligase domain^9–11^, and hormone receptors act as transcription factors that direct epithelial differentiation in breast tissues^12–15^. *BRCA1* deficiency accordingly produces aberrant differentiation in the mammary epithelium, implicating non-genetic mechanisms in tumor initiation. The events that convert this altered differentiation state into malignancy remain undefined.

*BRCA1* carriers predominantly develop triple-negative breast cancer (TNBC), defined by the absence of the estrogen and progesterone receptors and HER2. TNBC presents at a younger age, is more often metastatic at diagnosis and relapses more frequently than other breast cancers^16–18^. Treatment beyond surgery, chemotherapy and radiation is limited to PARP inhibitors and immune checkpoint blockade, and a substantial fraction of patients fail to respond or acquire resistance for reasons that mutational profiling has not resolved^19–25^. This variability reflects in part the fact that TNBC comprises several transcriptionally distinct diseases. Transcriptome-wide profiling resolves four subtypes, basal-like 1 (BL1), basal-like 2 (BL2), mesenchymal (M) and luminal androgen receptor (LAR), which differ in prognosis and in response to both drug classes^26–31^. TNBC-M is associated with epithelial-mesenchymal transition and stem-like transcriptional programs and with a high frequency of distant metastasis^26,27,32^. The determinants of subtype identity are unknown, and subtype is not used to stratify *BRCA1* carriers for treatment.

Genetically engineered mouse models have been the principal tool for investigating mechanisms of cancer initiation in *BRCA1* carriers. Because *TP53* is the most frequent co-mutation in *BRCA1*-driven breast cancer, most mouse models combine conditional deletion of *Brca1* with deletion or heterozygosity of *p53*. Tumors resembling human TNBC were shown to arise from at least two epithelial cell types: *Blg-Cre*-driven *Brca1* deletion^33^ generates tumors resembling invasive ductal carcinoma and are thought to be derived from luminal progenitor cells, whereas *Krt14-Cre*-driven deletion in basal cells generates tumors with a distinct histopathology resembling metaplastic cancer that is not commonly observed in *BRCA1* carriers^34^. The *Blg-Cre Brca1^f/f^p53^+/-^* model^33^ has therefore since dominated the field, and single-cell analyses have established that aberrant luminal differentiation and epithelial-to-mesenchymal transition precede tumor formation in this model^35,36^. However, this model fails to recapitulate the spectrum of breast cancer phenotypes observed in breast cancer patients.

To determine what specifies subtype in *Brca1*-associated disease, we characterized a less-studied model, *Wap-Cre Brca1^f^*^10*/f10*^*p53^f^*^5*&6/f5&6*^, in which loss of *Brca1* drives the ductal hyperplasia, excessive side branching and precocious alveologenesis of the mutant gland^37^, and whose cell of origin has not been established. Because *Wap* is expressed in secretory luminal cells, we anticipated a second LumSec-origin model. Combining single-cell transcriptomics, single-cell genotyping, lineage-restricted transplantation, spatial proteomics and integration with developmental and human breast cell atlases, we instead find that these tumors originate in Bmyo cells and correspond to human TNBC-M, whereas LumSec-derived *Blg-Cre* tumors correspond to TNBC-BL1. Cell of origin therefore shapes TNBC subtype, and Bmyo-derived disease is not restricted to metaplastic breast cancer.

## Results

### Aberrant epithelial differentiation in mammary tissues from Wap-Cre Brca1/p53 mice

We investigated the mechanisms of cancer initiation associated with BRCA1 deficiency by single-cell analysis of tissues from *Wap-Cre Brca1/p53* mutant mice (*Wap-Cre;Brca1^f10/f10^p53^f5&6/f5&6^*), which harbor loxP sites flanking exon 10 of *Brca1* and exons 5 and 6 of *p53* (Fig. 1a). The mammary glands of mutant mice exhibit ductal hyperplasia, excessive side branching and precocious alveolar formation, and the animals develop spontaneous tumors at approximately 6 months of age (Fig. 1b and Supplementary Table 1). To capture early tumorigenic events, we collected mammary tissues from 4-month-old mutant females that showed no visible tumor lesions, together with wild-type littermates for comparison. We also collected tumors ranging from 1 to 25 mm in diameter to investigate changes associated with tumor progression (Fig. 1c). Tissues were digested to single-cell suspensions and multiplexed for droplet-based single-cell RNA sequencing (scRNA-seq). Libraries were processed to remove low-quality cells, doublets and ambient RNA, yielding a dataset of 62,195 cells for downstream analysis (Fig. 1e and Supplementary Fig. 1a-e).

**Figure 1.**
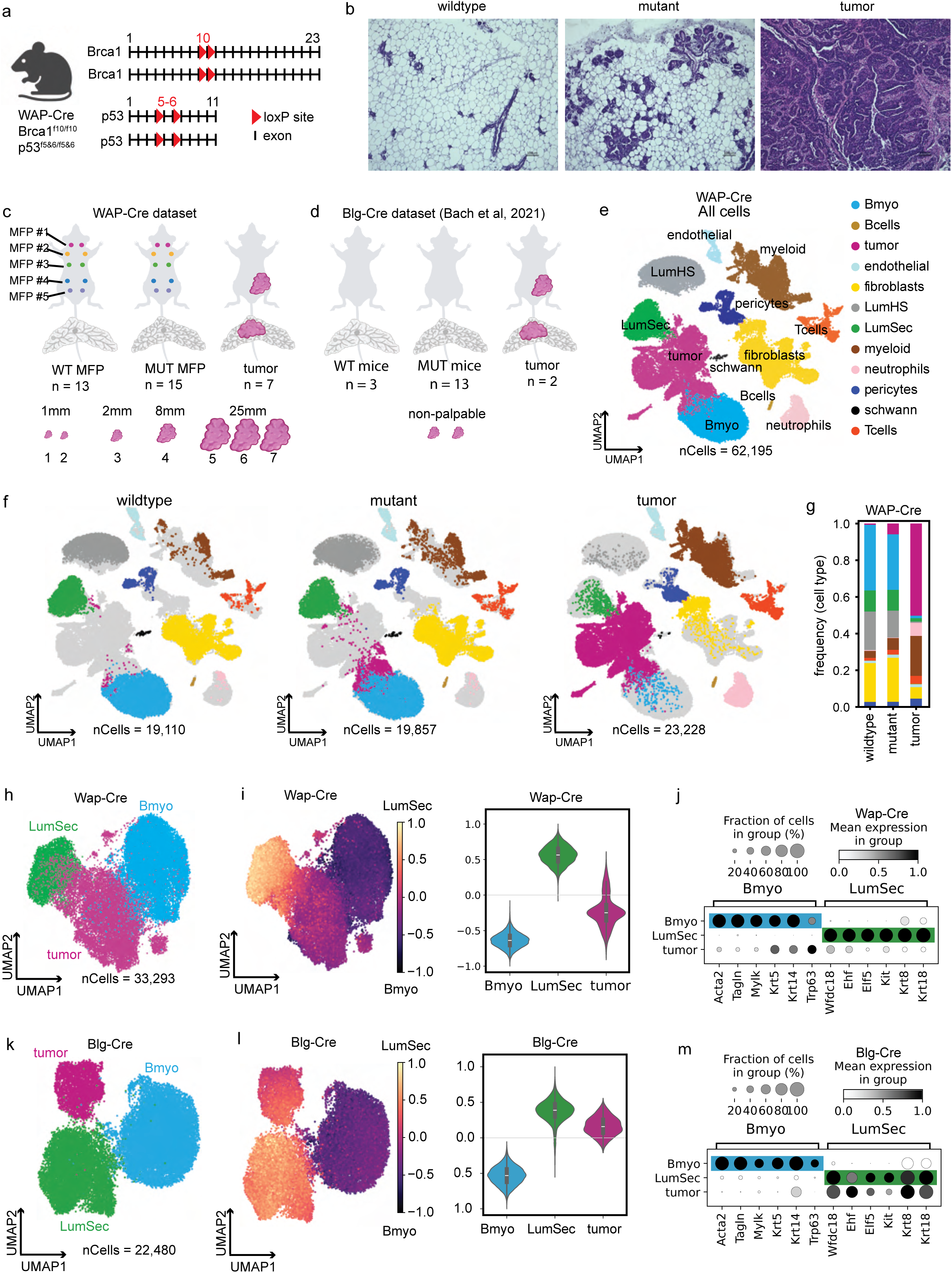
*Wap-Cre* and *Blg-Cre* mouse models exhibit transcriptomically distinct *Brca1* mammary tumors. **a,** Schematic of the *Wap-Cre Brca1* mouse model genetics. **b,** Representative H&E-stained mammary sections from wild-type (4 months), pre-tumor mutant (4 months) and tumor-bearing mutant (6 months) mice. Scale bar, 100 μm. Data are representative of *5* independent experiments. **c,d,** Experimental design for the current *Wap-Cre* dataset **(c)** and a previously published *Blg-Cre* dataset **(d;).** In c, individual mammary glands (1–5) from wild-type (*n* = 3 mice; 13 glands total) and mutant (*n* = 3 mice; 15 glands total) were separately barcoded and multiplexed with individual tumors (*n* = 7; 1–25 mm diameter). In d, mammary glands (1–5) were pooled within each mouse for wild-type (*n* = 3 mice) and mutant (*n* = 13 mice) cohorts, while non-palpable tumors (*n* = 2) were processed independently. **e,** UMAP projection of *Wap-Cre* mammary cells (*n* = 62,195 cells) passing quality control, colored by annotated cell type. **f,** UMAP plots partitioned by genotype and disease stage (wild-type, mutant and tumor). **g,** Relative cell-type frequencies across the conditions shown in f. **h–j,** Sub-analysis of Bmyo, LumSec and tumor cells in the *Wap-Cre* model including UMAP re-embedding **(h),** Bmyo–LumSec similarity scores **(i)** and expression of canonical Bmyo and LumSec markers **(j).** In i, the composite similarity score represents the additive enrichment of Bmyo and LumSec gene signatures, rescaled from –1 (Bmyo-like) to +1 (LumSec-like). Center line, median; box limits, upper and lower quartiles; whiskers, 1.5x the interquartile range. In j, dot size represents the percentage of cells expressing the marker; color intensity indicates average expression. **k–m,** UMAP re-embedding **(k),** Bmyo–LumSec similarity scores **(l)** and marker expression **(m)** for the *Blg-Cre* model, as described for h–j.

We identified 12 cell types -- four stromal, four immune and four epithelial -- using unbiased Leiden clustering and marker gene analysis (Fig. 1e, Extended Data Fig. 1c and Supplementary Table 2). Stromal populations included Schwann cells (*Plp1*), endothelial cells (*Pecam1*), pericytes (*Rgs5*) and fibroblasts (*Lum*). B cells (*Igkc*), T cells (*Cd3e*), neutrophils (*Cxcl2*) and myeloid cells (*Ctss*) were identified in the immune compartment. We found striking differences in the frequencies of these cell types among the three conditions -- wild-type, mutant and tumor (Fig. 1f,g and Extended Data Fig. 1d). Tumor tissues were enriched for myeloid cells, T cells and neutrophils. Whereas neutrophil expansion became evident only in tumors, myeloid and T cell expansion was already apparent in some mutant glands. T cells decreased in frequency with increasing tumor size. Fibroblasts and pericytes also changed with tumor progression, shifting markedly in UMAP space to become more similar to one another and showing substantial changes in gene expression (Fig. 1f). Differential abundance testing across cellular neighborhoods^38^ confirmed these shifts (Extended Data Fig. 1e).

We identified three canonical cell types in the epithelial compartment: basal-myoepithelial (Bmyo; *Krt14*, *Acta2*), luminal secretory adaptive precursor^39,40^ (LumSec; *Wfdc18*, *Kit*) and luminal hormone-sensing (LumHS; *Esr1*, *Pgr*) cells (Fig. 1e and Extended Data Fig. 1c). We also observed an intermediate population that bridged the Bmyo and LumSec populations in UMAP space. These cells were enriched in tumor samples and scored high for aneuploidy by CopyKAT^41^, indicating that they are tumor cells (Extended Data Fig. 1a,b). Importantly, a minority of cells with this phenotype were consistently observed in mutant tissues (approximately 10% of epithelial cells) but were nearly undetectable in wild-type glands (<1%), suggesting that they represent early transformation events.

Analysis of gene expression in the tumor cell cluster showed a lack of *Esr1*, *Pgr* and *Erbb2*, indicating that these tumors are triple-negative. The tumor cells expressed markers of both LumSec (*Wfdc18*) and Bmyo (*Krt14*) cells and displayed a much broader expression of Bmyo genes (*Krt5*, *Krt42*, *Trp63*, *Lgals7*, *Nkain3*) than has been reported in the *Blg-Cre Brca1/p53* model (*Blg-Cre;Brca1^f21–23/f21-23^p53^f/+^*). Tumors in *Blg-Cre* mice have been reported to develop as TNBC via aberrant differentiation of LumSec cells. To compare the two models directly, we reanalyzed publicly available scRNA-seq data from the *Blg-Cre* model^35^, which included wild-type, mutant and tumor samples (Fig. 1d). Using the same pipeline applied to our *Wap-Cre* dataset, we identified 12 main cell types, including three epithelial cell types and a tumor cell cluster that scored high for aneuploidy (Extended Data Fig. 1f-j and Supplementary Fig. 1h-j). Of note, tumor cells in the *Blg-Cre* model formed a discrete cluster that did not bridge Bmyo and LumSec cells (Fig. 1k and Extended Data Fig. 1f).

For higher-resolution analysis, we subset and re-clustered Bmyo, LumSec and tumor cells from both models (Fig. 1h,k and Supplementary Fig. 1f,k). This showed that tumor cells from the two models occupy different positions relative to Bmyo and LumSec cells in UMAP space. Scoring cells along a composite Bmyo-LumSec differentiation axis showed that tumor cells from the *Wap-Cre* model scored toward the Bmyo end, whereas tumor cells from the *Blg-Cre* model scored toward the LumSec end (Fig. 1i,l). Consistent with this, *Blg-Cre* tumor cells expressed much lower levels of the canonical basal markers *Krt5* and *Trp63*, and higher levels of the luminal markers *Wfdc18*, *Krt8,* and *Ehf*, than *Wap-Cre* tumor cells (Fig. 1j,m). These findings suggest that tumor progression in the two models follows distinct differentiation trajectories, raising questions about their relative relevance to disease in human *BRCA1* mutation carriers.

### Tumors from Wap-Cre Brca1/p53 mice recapitulate the mesenchymal subtype of TNBC in BRCA1 carriers

We next compared gene expression in the *Wap-Cre* and *Blg-Cre* models with TNBC tumors from *BRCA1* mutation carriers (Fig. 2a). Four subtypes of TNBC have been identified on the basis of transcriptome-wide gene expression: basal-like 1 (BL1), basal-like 2 (BL2), mesenchymal (M) and luminal androgen receptor (LAR)^28^. Also, a small fraction of tumors cannot be classified owing to insufficient similarity to any subtype and are designated unclassified (UNC)^28^. The four subtypes engage distinct biological programs (Fig. 2b) and differ in prognosis and therapeutic susceptibility. TNBC-M in particular is enriched for epithelial-mesenchymal transition and stem-like programs and is the subtype most frequently associated with distant metastasis.^27,32^

**Figure 2.**
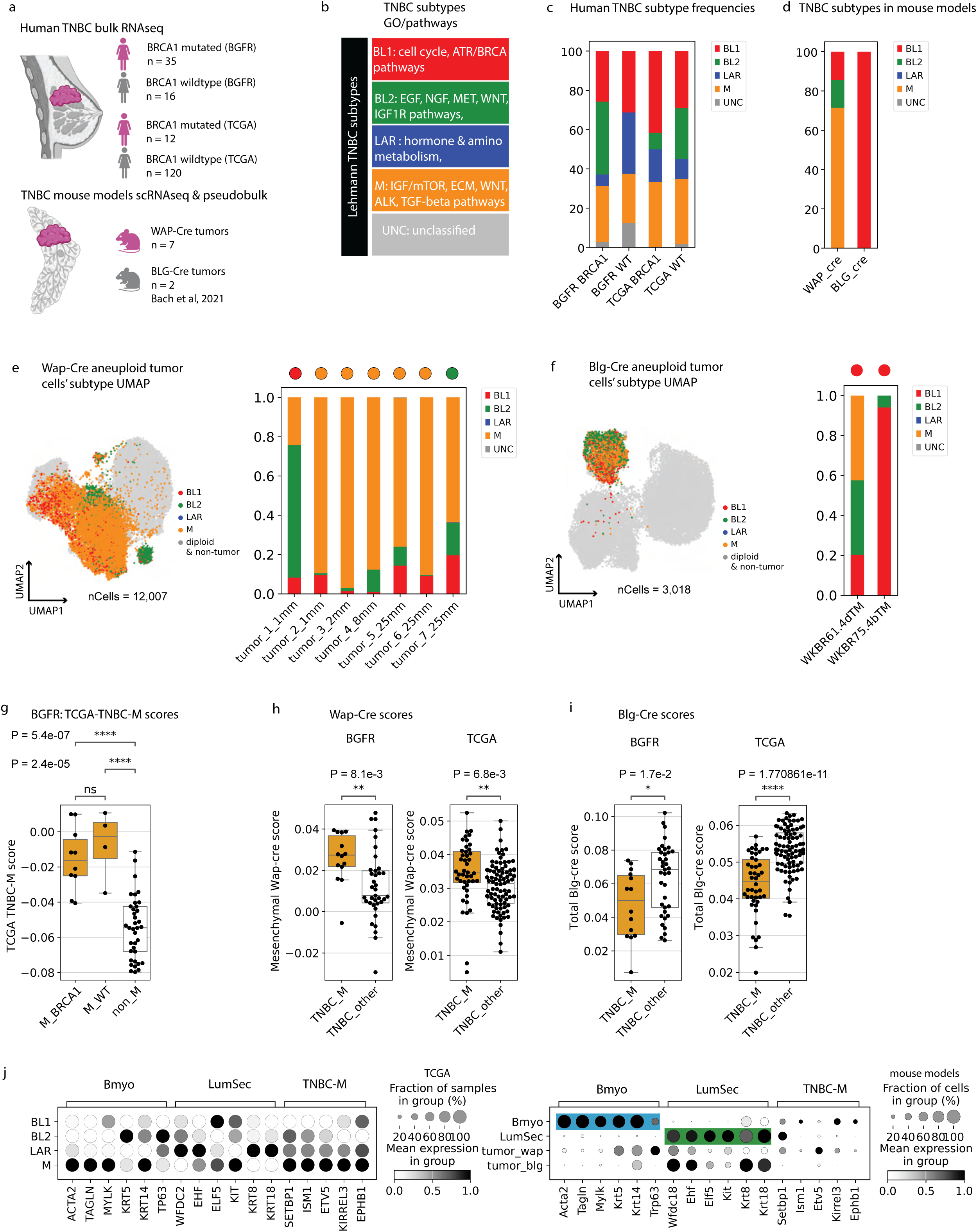
*Wap-Cre Brca1*-deficient tumors recapitulate the transcriptional program of human mesenchymal TNBC. **a,** Schematic of the experimental design comparing human TNBC clinical samples with *Wap-Cre* and *Blg-Cre Brca1*-deficient mouse models. Clinical samples comprise four groups from two independent cohorts: Basser-Gray Foundation Repository (BGFR) *BRCA1* carriers (*n* = 35) and non-carriers (*n* = 16), and TCGA *BRCA1* carriers (*n* = 12) and non-carriers (*n* = 120). **b,** Gene Ontology (GO) and pathway enrichment terms for TNBC subtypes as defined by Lehmann et al. (2011). **c,d,** Relative distribution of TNBC subtypes in human clinical cohorts stratified by *BRCA1* status **(c;** *n* = 183 samples) and in *Wap-Cre* and *Blg-Cre* mouse models **(d;** *n* = 9 tumors). **e,f,** Single-cell TNBC subtyping in *Wap-Cre* **(e)** and *Blg-Cre* **(f)** tumors. UMAP projections (left) and stacked bar plots (right) show the relative proportions of subtyped cells (*n* = 15,025 cells), restricted to the aneuploid tumor cell population. Circles above each bar indicate the predicted TNBC subtype based on pseudo-bulk analysis shown in **d**. **g–i,** Box plots of enrichment scores for the human TNBC-M signature **(g)**, the *Wap-Cre* mesenchymal tumor signature **(h)** and the *Blg-Cre* tumor signature **(i)** in human clinical samples. Scoring in **g** is restricted to the BGFR cohort (*n* = 51), as the human TNBC-M signature was derived from TCGA; in **h** and **i**, the BGFR (*n* = 51) and TCGA (*n* = 132) cohorts are shown separately, as labeled. The *Wap-Cre* signature was derived from the 5 of 7 mouse tumors classified as mesenchymal in **d** and the *Blg-Cre* signature from all *Blg-Cre* tumor cells, in each case by comparison with wild-type epithelium (see **Methods**). Samples are stratified into mesenchymal (M) and non-mesenchymal subtypes throughout; in **g**, the M group is further sub-stratified by *BRCA1* mutation status (for **h** and **i**, see **Extended Data** Fig. 2f). Center line, median; box limits, upper and lower quartiles; whiskers, 1.5x the interquartile range. *P* values determined by one-way ANOVA followed by Tukey’s post-hoc test, performed separately within each cohort; cohorts were not compared. **j,** Dot plot showing expression of canonical basal-myoepithelial (Bmyo) and luminal-secretory (LumSec) markers, alongside TNBC-M marker genes derived via differential expression analysis of TCGA human TNBC subtypes (see **Methods** for gene list derivation and **Supplementary Table 3** for full gene list). Color intensity represents mean expression level; dot size represents the percentage of patient samples in TCGA subtypes expressing the gene (left) and percentage of cells expressing the gene in mouse datasets (right).

We determined whether *BRCA1* mutation carriers preferentially develop specific subtypes of TNBC. We analyzed bulk RNA-sequencing data collected from 51 patients with TNBC (Basser-Gray Foundation Repository, BGFR; 35 *BRCA1* carriers and 16 non-carriers, of which 17 samples were previously described^24^ and 34 are newly reported here), together with a TNBC cohort from The Cancer Genome Atlas (TCGA, *n* = 132; 12 *BRCA1* carriers and 120 non-carriers as determined previously^42^). Subtypes were assigned by determining the highest centroid correlation between each tumor and the established TNBC subtype centroids^43^. This showed that *BRCA1* carriers develop all TNBC subtypes, in roughly the same proportions as in sporadic TNBC (Fig. 2c and Supplementary Table 5). We then performed the same subtyping analysis on pseudo-bulked scRNA-seq profiles from the mouse models, and at single-cell resolution in the epithelium alone. We found that the two *Brca1* models recapitulate different TNBC subtypes. *Wap-Cre* tumors were predominantly assigned the mesenchymal (TNBC-M) subtype (5 of 7 tumors), whereas *Blg-Cre* tumors were assigned basal-like 1 (TNBC-BL1) (Fig. 2d and Supplementary Table 7). At single-cell resolution, a surprising degree of intra-tumor subtype heterogeneity was evident in both models (Fig. 2e,f). Although the dominant subtype in the cancer epithelium did not always match the tissue-level pseudo-bulk assignment, there was substantial concordance, and the *Wap-Cre* model showed a higher frequency of cells resembling the TNBC-M subtype (Fig. 2e,f).

We next investigated whether any differences exist between BRCA1-associated and sporadic TNBC-M, and whether tumor cells from the *Wap-Cre* tumors classified as mesenchymal resemble human TNBC-M at the level of gene expression. We derived a human TNBC-M signature by contrasting mesenchymal TNBCs against non-mesenchymal tumors in the TCGA breast cancer cohort and used the gene signature for module scoring (Supplementary Table 3). Applied to the 51 BGFR TNBC samples, this signature behaved as expected, being enriched in tumors classified as TNBC-M (Fig. 2g) with no differences based on BRCA1 mutation status. We then generated transcriptomic signatures from the five mesenchymal *Wap-Cre* tumors and from both *Blg-Cre* tumors by contrasting tumor cells against wild-type epithelial cells, and used them for module scoring of the BGFR and TCGA tumors. Because the contributing tumors were selected by the classifier, this analysis supports the subtype assignments rather than testing them independently. The contrast is against normal epithelium rather than between subtypes, so the classifier determined which tumors contributed to each signature but not which genes defined it. To ensure that the tumor epithelium and not the stroma explained the TNBC-M designation in the mouse tumors, the mouse signatures were derived from the epithelial compartment only. The *Wap-Cre* signature scored highest in TNBC-M tumors in both cohorts (BGFR: *P* = 0.0081; TCGA: *P* = 0.0068), whereas the *Blg-Cre* signature, derived in the same way from tumors assigned to TNBC-BL1, did not (Fig. 2h,i).

We next examined TNBC-M gene expression in greater detail across the epithelium, focusing on the top 125 differentially expressed genes in the derived signature. We found that *Wap-Cre* tumors express higher levels of TNBC-M genes such as *Setbp1, Etv5, Kirrel3* than *Blg-Cre* tumors (Fig. 2j and Extended Data Fig. 2b). *Wap-Cre* tumors and human TNBC-M tumors both express higher levels of canonical Bmyo genes and lower levels of LumSec genes (Fig. 2j and Extended Data Fig. 2b). Together, these results show that the two widely used *Brca1* GEMMs recapitulate different subtypes of human TNBC, both of which occur in *BRCA1* mutation carriers. Furthermore, TNBC-M appears to be associated with more basal differentiation, and TNBC-BL1 with more LumSec differentiation.

### Aberrant basal cell differentiation in pre-malignant tissues from Wap-Cre Brca1/p53 mice

We investigated the ontogeny of disease initiation in the *Wap-Cre* and *Blg-Cre* models by further analysis of mammary glands from mutant mice before overt tumor development. Mutant glands from both models contained aneuploid cells, indicating that early transformation events are already present at this stage (Fig. 3a-h). To evaluate differentiation in these cells, we performed subset analysis of the aneuploid cells together with Bmyo and LumSec cells from wild-type and mutant mice (Fig. 3b,c,f,g). In the *Wap-Cre* model, aneuploid cells were heterogeneous, clustering with both Bmyo and LumSec cells but also forming a separate cluster of tumor-like cells (Fig. 3b-d). This tumor-like cluster contained the majority of aneuploid cells and clustered more closely with Bmyo cells. Canonical marker gene analysis showed that tumor-like cells expressed basal genes and TNBC-M genes (Fig. 3i), and they scored toward the Bmyo end of the Bmyo-LumSec axis (Fig. 3j). In contrast, in the *Blg-Cre* model the majority of aneuploid cells clustered with LumSec cells, expressed canonical LumSec markers, scored toward the LumSec end of the axis and did not express TNBC-M genes (Fig. 3f-h,k,l). Tumor initiation in the *Wap-Cre* model is therefore associated with aberrant basal cell differentiation, whereas in the *Blg-Cre* model it is restricted to aberrant luminal cell differentiation.

**Figure 3.**
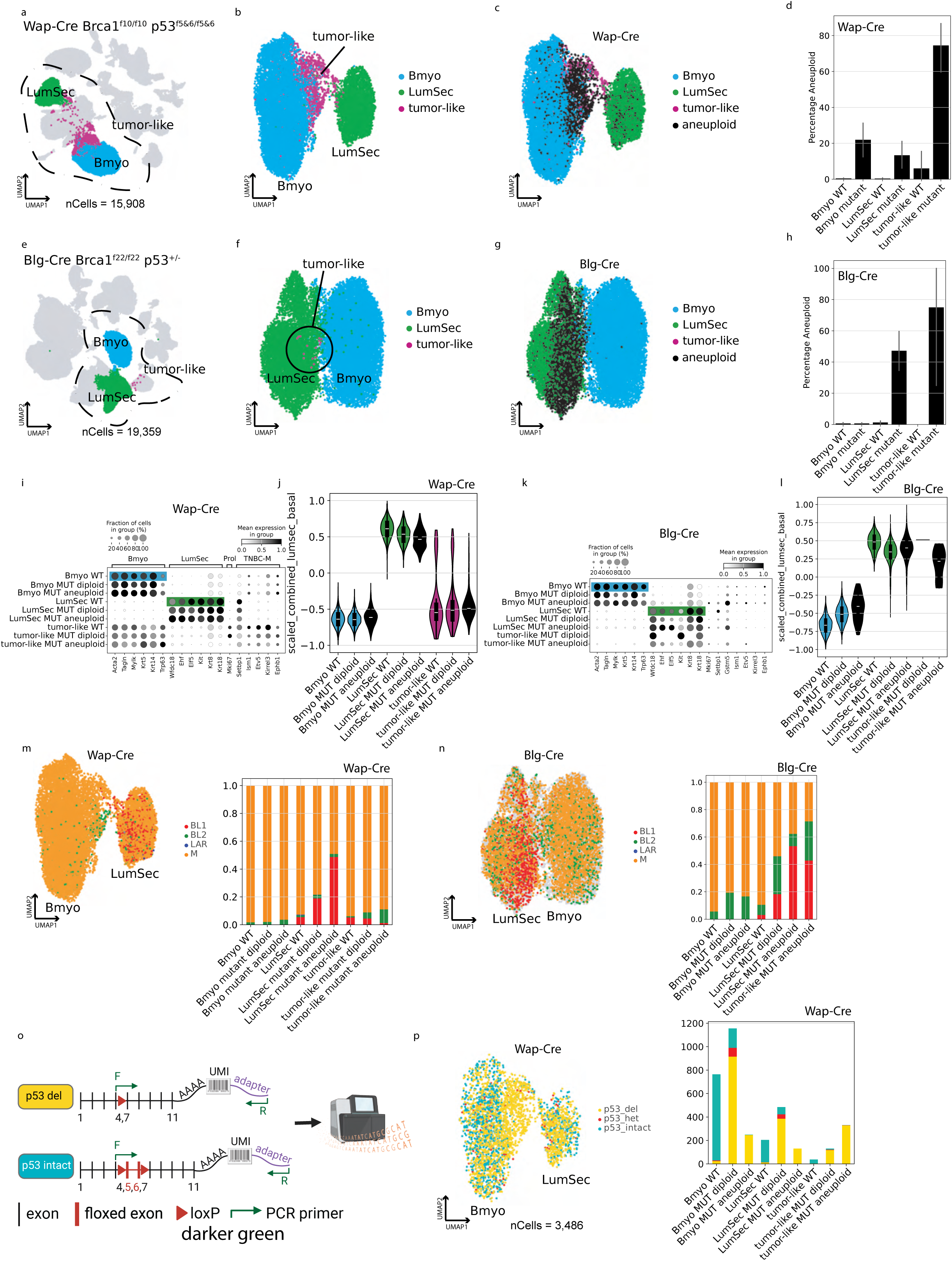
*Brca1*-deficient mammary models exhibit early transcriptional divergence and distinct cells-of-origin. **a,** UMAP visualization of all cells in the *Wap-Cre* model. Wild-type and mutant epithelial cells are colored by cell type; epithelial clusters are encircled. **b,c,** Re-embedding of the epithelial populations from a. Cells are colored by cell type **(b)** or labeled by ploidy if aneuploid and by cell type if diploid **(c)**. **d,** Mean percentage of aneuploid cells within each epithelial cell type, stratified by condition (*n* = 13 wild-type vs. *n* = 15 mutant). Data represent mean ± s.e.m. **e–h,** Analogous characterization for the *Blg-Cre* model, including global UMAP **(e),** epithelial re-embedding **(f)** and identification of aneuploid cells (*n* = 3 wild-type vs. *n* = 13 mutant) **(g,h)** as described in c,d. **i,** Dot plot of *Wap-Cre* data from **c** showing expression of canonical Bmyo, LumSec and TNBC-M marker genes (defined in Fig. 2 and **Methods**). Data are stratified by cell type and condition, with mutant cells further sub-stratified by ploidy. **j,** Violin plots with overlaid box plots displaying Bmyo–LumSec composite scores for groups defined in **i.** Box plot center line, median; box limits, upper and lower quartiles; whiskers, 1.5x interquartile range. **k,l,** Dot plot and violin plot analyses for the *Blg-Cre* model, as described in i,j. **m,n,** Single-cell TNBC subtype-like assignment visualized on UMAPs (left) and stacked bar plots (right) for *Wap-Cre* **(m)** and *Blg-Cre* **(n)** models. **o,** Schematic of the single-cell genotyping approach. A customized MAESTER-based approach captures *Trp53* transcripts, enabling discrimination between floxed and recombined alleles based on exon 5–6 retention. **p,** Single-cell genotyping results visualized on a UMAP (left) and via non-normalized stacked bar plots (right). Bar heights represent raw cell counts; colors indicate genotype proportions across cell types, conditions and ploidy states.

We next performed single-cell subtyping analysis of aneuploid cells from each model to look for correlations between differentiation state and subtype assignment. The vast majority of *Wap-Cre* tumor-like cells were assigned the M subtype, whereas the predominant subtype among *Blg-Cre* tumor-like cells was BL1 (Fig. 3m,n). The non-malignant epithelium showed the same association. Mutant LumSec cells in both models were enriched for BL1-like assignment, further supporting a LumSec origin for *Blg-Cre* tumors, and mutant Bmyo cells in both models were enriched for M-like assignment, further supporting a Bmyo origin for *Wap-Cre* tumors. The subtype composition of the aneuploid compartment in each model therefore mirrors that of its putative cell of origin.

To determine whether the basal-enriched disease phenotype observed in the *Wap-Cre* model occurs because the Cre is preferentially expressed in basal cells, we performed single-cell genotyping to assess which cell types display gene deletion. Because the *Brca1* locus is too large for efficient multiplexed PCR amplification, we adapted the MAESTER variant enrichment framework^44^ to capture *Trp53* transcripts, which are recombined by the same *Wap*-driven Cre (Fig. 3o). Remarkably, we found that both Bmyo and LumSec cells in *Wap-Cre* mutant mice carry recombined alleles (Fig. 3p, Extended Data Fig. 3d and Supplementary Fig. 3a), confirming that both cell types accrue Cre-driven mutations and that restricted recombination does not account for the basal phenotype of the disease. In contrast, analysis of the published *Blg-Cre* dataset by CNV inference indicates that only LumSec cells transform in that model (Fig. 3h). Together, these data suggest that aberrant basal cell differentiation yields the TNBC-M subtype, whereas aberrant luminal cell differentiation yields TNBC-BL1.

### Basal-derived tumors reproduce the phenotype of spontaneous *Wap-Cre* tumors

To functionally investigate the cell-of-origin of disease in the *Wap-Cre model,* we performed transplants of basal and luminal cells from mutant *Wap-Cre* animals prior to tumor onset (Fig. 4a). Basal and luminal cells were sorted by flow cytometry using a standard marker panel (Lin, EpCAM, CD49f) from 4-month-old mutant mice lacking overt tumor lesions (Fig. 4b). The sorted cells were combined with stromal cells from the same donors and transplanted into the cleared fat pads of immunodeficient NSG recipients. Eight recipients received contralateral implants of basal and luminal cells, and one received basal cells bilaterally. A sample of cells from each fraction was also analyzed by scRNA-seq. Clustering analysis confirmed successful fractionation with minimal cross-contamination between basal and luminal identities, and both fractions contained comparable amounts of aneuploid cells (Fig. 4c-e and Extended Data Fig. 4b).

**Figure 4.**
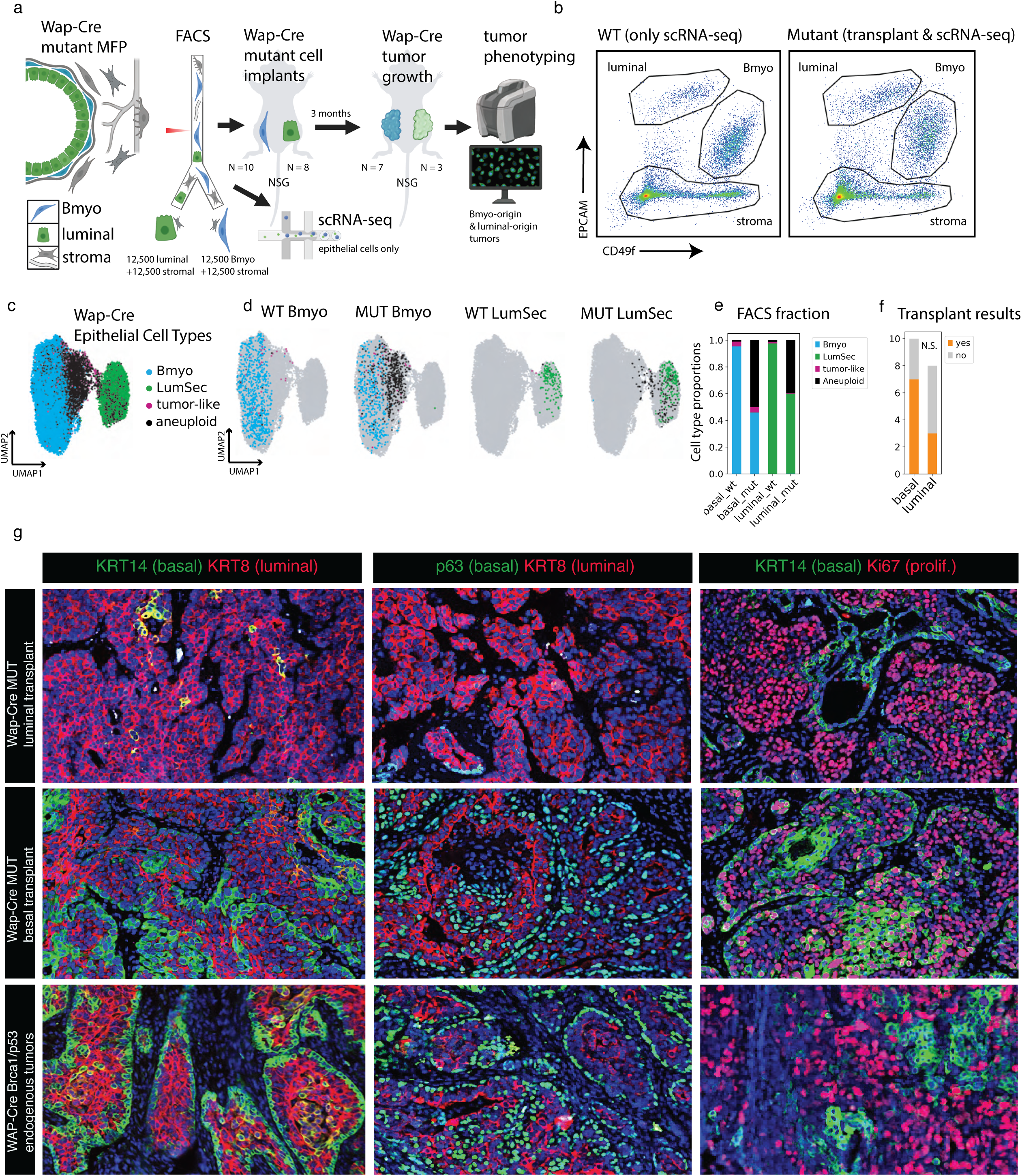
*Wap-Cre* Bmyo cell fraction harbors transplantable tumor-initiating cells with a capacity to produce tumors that recapitulate spontaneous phenotypes. **a,** Experimental workflow for the isolation and orthotopic transplantation of specific mammary epithelial subsets. Basal and luminal populations were FACS-purified from mutant *Wap-Cre* donors and implanted into the cleared mammary fat pads of female NSG (NOD-*scid* Il2rg^null^) recipient mice (at *n* = 25,000 cells per injection). **b,** Representative FACS plots and gating strategy for the isolation of mammary epithelial cell (MEC) subsets from wild-type (left) and mutant (right) donor mice. MECs were defined as Lin^-^ (Cd45^-^) and further partitioned into basal-myoepithelial (Epcam^mid^ CD49f^high^) and luminal (Epcam^high^ CD49f^mid^) populations. **c,** UMAP projection of epithelial cells from transplanted cohorts, labeled by ploidy status (if aneuploid) or by cell type (if diploid). **d,** UMAP plots (as in c) partitioned by genotype and the specific FACS gate of origin, demonstrating the transcriptomic purity of the sorted populations. **e,** Relative cell-type frequencies across the experimental conditions and gates shown in **d,** confirming minimal cross-contamination between basal and luminal identities. **f,** Frequencies of tumor formation in NSG recipients following the implantation of sorted basal or luminal mutant *Wap-Cre* cells (*n* = 8 injection sites in luminal group, *n* = 10 in basal group). Differences in tumor formation rates were not statistically significant (two-sided Fisher’s exact test, ***P*** = 0.34). **g,** Representative immunostained sections of tumors derived from transplanted pre-tumor mutant cells (top, middle) and spontaneously formed *Wap-Cre* tumors (bottom). Scale bar, 100 μm. Data are representative of *n* = 10 independent transplant-derived tumors (seven basal-derived and three luminal-derived) and *n* = 3 spontaneous tumors, with similar results across *n* = 2 independent experiments.

Basal and luminal cells produced tumors at statistically indistinguishable rates (7 of 10 versus 3 of 8 injection sites; two-sided Fisher’s exact test, *P* = 0.34; Fig. 4f), and tumors grew to comparable size, as determined by tumor mass at the end of the experiment (Extended Data Fig. 4f and Supplementary Table 4). Immunofluorescence analysis, however, showed that the tumors produced by basal and luminal cells were remarkably distinct (Fig. 4g). All seven tumors arising from the basal cell fraction contained basal cells (Krt14^+^ Trp63^+^), whereas none of the three tumors from the luminal cell fraction did. Tumors from both fractions contained luminal cells (Krt8^+^), as well as hybrid cells expressing markers of both the basal and luminal lineages. Spontaneous tumors generated in intact *Wap-Cre* animals showed a similar expression pattern as the tumors generated from basal cell transplants. Tumor formation therefore did not distinguish the two fractions, but tumor phenotype did: only basal-derived tumors reproduced the cellular composition of spontaneous tumors, supporting a basal cell of origin for TNBC-M in the *Wap-Cre* model (Fig. 4g). Consistent with both fractions being tumorigenic, the remaining 2 of 7 spontaneous *Wap-Cre* tumors were not assigned the mesenchymal subtype (Fig. 2d), as would be expected if a minority arise from luminal cells, which in the *Blg-Cre* model give rise to TNBC-BL1.

### Aberrant basal cell differentiation recapitulates fetal mammary development

We investigated whether the altered differentiation observed in the *Wap-Cre* model is entirely aberrant or instead resembles a specialized trajectory that appears during a particular stage of mammary development. The mammary gland is a highly dynamic organ that undergoes extensive tissue remodeling throughout the life cycle. We compared differentiation in the *Wap-Cre* and *Blg-Cre* models with a comprehensive developmental reference dataset (Dev-atlas) containing cells collected during fetal development, puberty, adult homeostasis, gestation, lactation and involution (Fig. 5a)^45^. The dataset also includes cells collected from fetal skin, the embryonic origin of the mammary gland, and cells microdissected from ducts and terminal end buds of pubescent glands.

**Figure 5.**
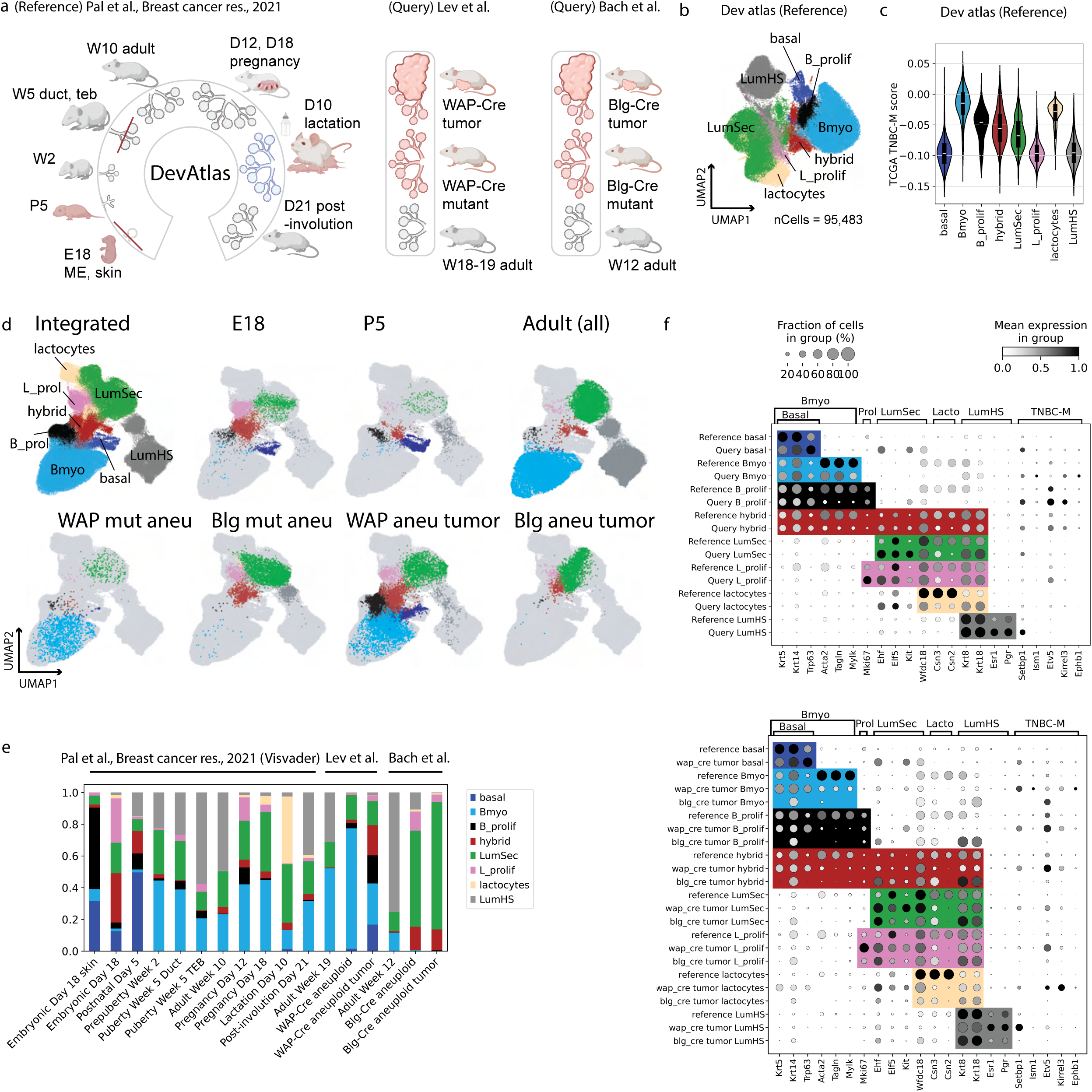
Integration with a developmental mammary atlas identifies the cellular ontogeny of *Brca1*-deficient pre-tumor and tumor populations. **a,** Schematic of the experimental design for the transcriptomic integration of a developmental mouse mammary epithelial cell atlas (Dev-atlas) with *Wap-Cre* and *Blg-Cre Brca1*-deficient datasets. **b,** UMAP projection of the Dev-atlas reference (*n* = 95,483 cells), reprocessed using the computational pipeline described in Fig. 1e and re-annotated with scANVI (see Methods). **c,** Violin plots with overlaid box plots displaying human TNBC-M signature enrichment scores across Dev-atlas-defined cell populations. Center line, median; box limits, upper and lower quartiles; whiskers, 1.5x the interquartile range. **d,** UMAP projections of the joint latent space generated by scANVI integration of the Dev-atlas and the *Brca1*-deficient datasets (*n* = 124,145 cells). Cells are partitioned by selected developmental stages and disease states. **e,** Relative cell-type frequencies within the integrated dataset, illustrating the distribution of Dev-atlas-defined populations identified through joint scANVI co-clustering across the samples defined in a. **f,** Dot plots showing expression of canonical markers and derived TNBC-M marker genes (defined in Fig. 2 and **Methods**). Data are stratified by Dev-atlas-defined cell populations and sub-stratified by integration batch (top) and selected biological conditions (bottom). Color intensity represents mean expression level; dot size represents the percentage of cells expressing the gene.

We re-analyzed the Dev-atlas reference dataset with the same pipeline used for adult tissues and observed far greater epithelial diversity than we observed in adult glands. Clustering and marker gene analysis identified eight epithelial cell states across the developmental stages (Fig. 5b,f and Extended Data Fig. 5a). As expected, adult glands were almost exclusively composed of the three main cell types: Bmyo, LumSec and LumHS. Specialized cell states were found during pregnancy, lactation and fetal development. Pregnant and lactating glands contained milk-producing alveolar cells (lactocytes^39^; *Csn2*, *Wfdc18*). Fetal skin and fetal mammary gland contained a unique basal cell type that expresses canonical basal cell markers but lacks myoepithelial markers (basal; *Krt5^+^*, *Krt14^+^*, *Trp63^+^*, *Acta2^-^*, *Tagln^-^*, *Mylk^-^*). Previous work suggests that these are fetal stem cells that give rise to basal and luminal cells^45–48^. We also found hybrid cells expressing markers of both basal and luminal cells (hybrid; *Krt14*, *Acta2*, *Krt8*, *Wfdc18*), which prior work indicates represent intermediate or transitional cells^45,46,48^. Hybrid cells were most abundant in fetal development, but small frequencies were also found at most stages. Proliferating populations of basal and luminal cells were likewise most abundant during fetal development and pregnancy, when the gland undergoes extensive growth. Module scoring of each cell state for the human TNBC-M signature gave generally negative scores, indicating dissimilarity to TNBC-M tumors, but Bmyo cells and lactocytes had the least negative scores (Fig. 5c).

We integrated and co-clustered the *Wap-Cre* and *Blg-Cre* datasets with the Dev-atlas reference. Integration was effective, as determined by the percentage of label agreement before and after integration and co-clustering, and the joint latent space showed no residual batch structure (Fig. 5d and Supplementary Fig. 5b,c). This analysis revealed more striking differences in epithelial differentiation between the *Wap-Cre* and *Blg-Cre* models. Aneuploid cells from the *Blg-Cre* model clustered predominantly with LumSec cells, in addition to low frequencies of proliferating LumSec and hybrid cells, suggesting that disease in this model is restricted to the luminal lineage (Fig. 5d,e and Extended Data Fig. 5b). In contrast, aneuploid tumor cells from the *Wap-Cre* model showed a greater diversification of cell states, with a large expansion of the fetal basal cell type, as well as hybrid and proliferative cells in addition to cells resembling adult Bmyo and LumSec cells (Fig. 5d,e and Extended Data Fig. 5b,c). Interestingly, analysis of aneuploid cells from the mutant gland of *Wap-Cre* mice (prior to tumor onset) showed primarily Bmyo and LumSec cells, with a low frequency of the hybrid or basal cell types, suggesting that disease initiation in these animals is more likely to occur through de-differentiation of Bmyo cells, than through expansion of a small population of residual hybrid or basal cells that persist beyond fetal development.

Examination of the seven tumors individually revealed an interesting progression relating to tumor size. The two smallest tumors (1 mm) resembled mutant glands, and were composed primarily of Bmyo cells and small frequencies of basal, hybrid, and LumSec cells (Extended Data Fig. 5b,c). The 2 and 8 mm tumor contained increasing frequencies of basal and proliferating Bmyo cells. Finally, the three 25 mm tumors showed a large expansion of hybrid cells, proliferating luminal cells and LumSec cells (Extended Data Fig. 5b,c). Tumor progression therefore appears to begin with expansion of aneuploid Bmyo-like cells, towards increasing differentiation and diversity of cell states, including the LumSec-like states.

Importantly, although tumor cells integrated with reference cells, we observed significant differences in gene expression between the healthy and tumor cells within each epithelial cell cluster (Fig. 5f). The most notable deviation was in the Bmyo cell state. Although tumor Bmyo cells clearly cluster with healthy Bmyo cells, suggesting they share some gene expression programs, the tumor Bmyo cells do not express the canonical myoepithelial markers *Acta2*, *Tagln* or *Mylk*. Loss of myoepithelial gene expression extended to the reference-defined hybrid state as well, in tumor cells of both *Brca1* models. This further indicates that disease initiation in the *Wap-Cre* model is associated with progressive de-differentiation of Bmyo cells into a basal cell state resembling that which is observed in fetal development.

We also performed single-cell TNBC subtyping on 5,587 fetal E18 cells from the reference atlas for comparison alongside adult cells. Fetal epithelium showed enrichment of TNBC-M-like cells across all cell states. Basal cells and lactocytes, although extremely rare in adult homeostasis, were also almost exclusively TNBC-M-like. Single-cell TNBC subtype distributions in reference-mapped cell states largely recapitulated the association we previously observed (Extended Data Fig. 5d,e). Bmyo-associated cell states were enriched for TNBC-M-like cells, whereas TNBC-BL1-like cells were restricted to the luminal lineage. The expanded set of reference-defined cell states in the adult mouse further showed that BL1 cells are more closely associated with proliferation (L_prol), while TNBC-LAR is associated with LumHS cells. In the tumor models, BL1 cells were detected across almost all *Blg-Cre* tumor cell states, while TNBC-M cells were dominant across *Wap-Cre* tumor cell states. These findings agree with our earlier assessment that the *Wap-Cre* and *Blg-Cre* models are fundamentally different, and further suggest that subtype-associated programs are largely retained as tumors differentiate.

### Aberrant basal cell differentiation localizes to regions of hyper-branching in mutant glands

To localize the aberrant differentiation observed in the *Wap-Cre* model in situ, we assayed wild-type glands, mutant glands, and small and large tumors using multiplexed proteomics (CODEX) (Fig. 6a,b). We used canonical epithelial markers that distinguish cells in the basal layer (Krt14^+^ SMA^+^ Krt8^-^) from cells in the luminal layer (Krt14^-^ SMA^-^ Krt8^+^) (Fig. 6c and Supplementary Table 6). We also compared proliferation in the two lineages using the proliferation marker MKI67. Wild-type epithelium showed the expected inner luminal layer and outer basal-myoepithelial layer (Fig. 6d). Of note, we were not able to detect cells with a hybrid (e.g., Krt14^+^ Krt8^+^) or fetal basal (e.g. Krt14^+^ SMA^-^) phenotype in wild-type glands (Fig. 6d). Interestingly, mutant glands contained regions of tissue that resembled wild-type glands (normal-like), but also displayed regions with extensive hyper-branching (hyperplastic) as reported previously (Fig. 6d and Supplementary Fig. 6a-f). Normal-like regions of mutant glands did not differ significantly from wild-type glands. However, we found that cells comprising the basal layer of hyperplastic regions did not express the myoepithelial marker SMA (Fig. 6d), consistent with the fetal basal cell phenotype observed in our single-cell RNA-sequencing dataset. Tumors were also highly enriched for such cells (Fig. 6e-g). Proliferation was infrequent in wild-type epithelium and occurred mostly in the luminal layer (Extended Data Fig. 6a,c). However, mutant glands contained a high frequency of proliferating cells, particularly within the hyperplastic regions, with the SMA^-^ basal cells proliferating most frequently. These data show that highly proliferative, aberrant basal epithelial differentiation localizes to regions of precious ductal branching, suggesting it is a key feature of disease onset in the *Wap-Cre* model.

**Figure 6.**
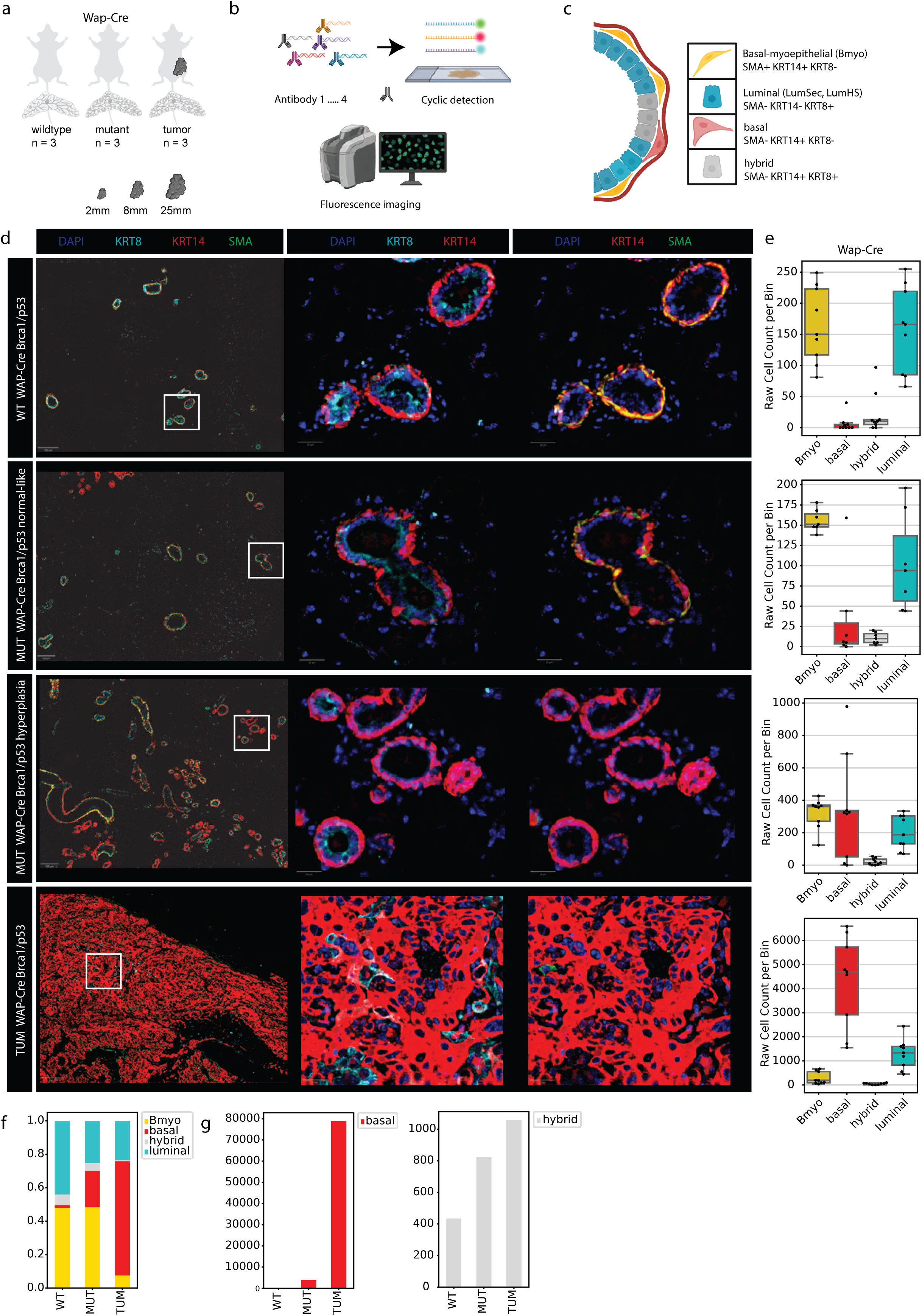
Spatial phenotyping by multiplex immunofluorescence (CODEX) validates early lineage alterations and cellular compositions in the *Wap-Cre* model. **a-c,** Experimental and technological workflow, showing the experimental design for CODEX multiplex immunofluorescence profiling across *Wap-Cre* mammary disease stages (**a**), and a diagram of iterative multicycles of fluorescent antibody staining and fluorophore inactivation (**b**), and a schematic of the mammary epithelial bilayer highlighting canonical cell lineage markers **(c)**. **d,** Representative multiplex immunostained sections of *Wap-Cre* mammary tissues across disease progression: wild-type (top), pre-tumor mutant normal-like (second row), mutant hyperplastic (third row) and tumor (bottom). Pseudocolored images depict key cell-type lineage markers as defined in **b**. Scale bars: overview images, 100 μm; inset images, 20 μm. Data are representative of n = 3 mice per condition, with similar results obtained across n = 3 independent experiments. **e,** Box plots with overlaid data points showing the density (cells per ROI) of segmented cell types across the experimental conditions described in **d**. Regions of interest (ROIs; 1 mm²) for normal-like and hyperplastic tissues were defined as described in Methods (all ROIs shown in Supplementary Fig. 6). Center line, median; box limits, upper and lower quartiles; whiskers, 1.5x the interquartile range. **f,** Stacked bar plot showing the relative cell-type proportions within each experimental condition. **g,** Bar plots displaying raw cell counts for basal (left) and hybrid (right) within each experimental condition.

### Metastatic lesions display a shift towards luminal differentiation while retaining a mesenchymal-like transcriptional program

Women with hereditary pathogenic variants in the *BRCA1* gene disproportionally develop aggressive TNBCs with a high propensity for the aggressive visceral and central nervous system (CNS) metastasis^49,50^. We investigated epithelial differentiation in experimental metastases generated following intracardiac injection of primary tumor cells from *Wap-Cre* mice, focusing on determining whether they demonstrate a shift towards aberrant basal or luminal differentiation. We dissociated a large primary tumor (tumor_5_25mm) and transplanted the cells intracardially into immunodeficient animals to generate multi-organ metastases (n = 6), or orthotopically (n = 2) to control for potential confounding effects of transplantation (Fig. 7a,b and Extended Data Fig. 7a,b). Lung, liver, brain, and mammary glands were harvested at endpoint and dissociated to single cell suspension. Metastatic cells were identified by flow cytometry based on strain specific allelic differences in MHCI expression between donor cells (H2-K^b^) and host cells (H2-K^d^) (Fig. 7c and Supplementary Fig. 7a,b). We profiled 4,806 cells from metastatic and orthotopic lesions, including an aliquot of the donor primary tumor carried through the identical enrichment workflow (757 cells) to control for any effect of MHC-based selection on recovered cell composition. Enriched and unenriched profiles of the donor tumor did not differ and were combined for subsequent analyses. All recipient mice grew orthotopic tumors or metastases, and several intracardiac recipients developed lesions in more than one organ. Take rates and metastatic burdens differed significantly among the peripheral organs, where some displayed very low frequencies of metastatic cells and others showed extensive metastatic burden.

**Figure 7.**
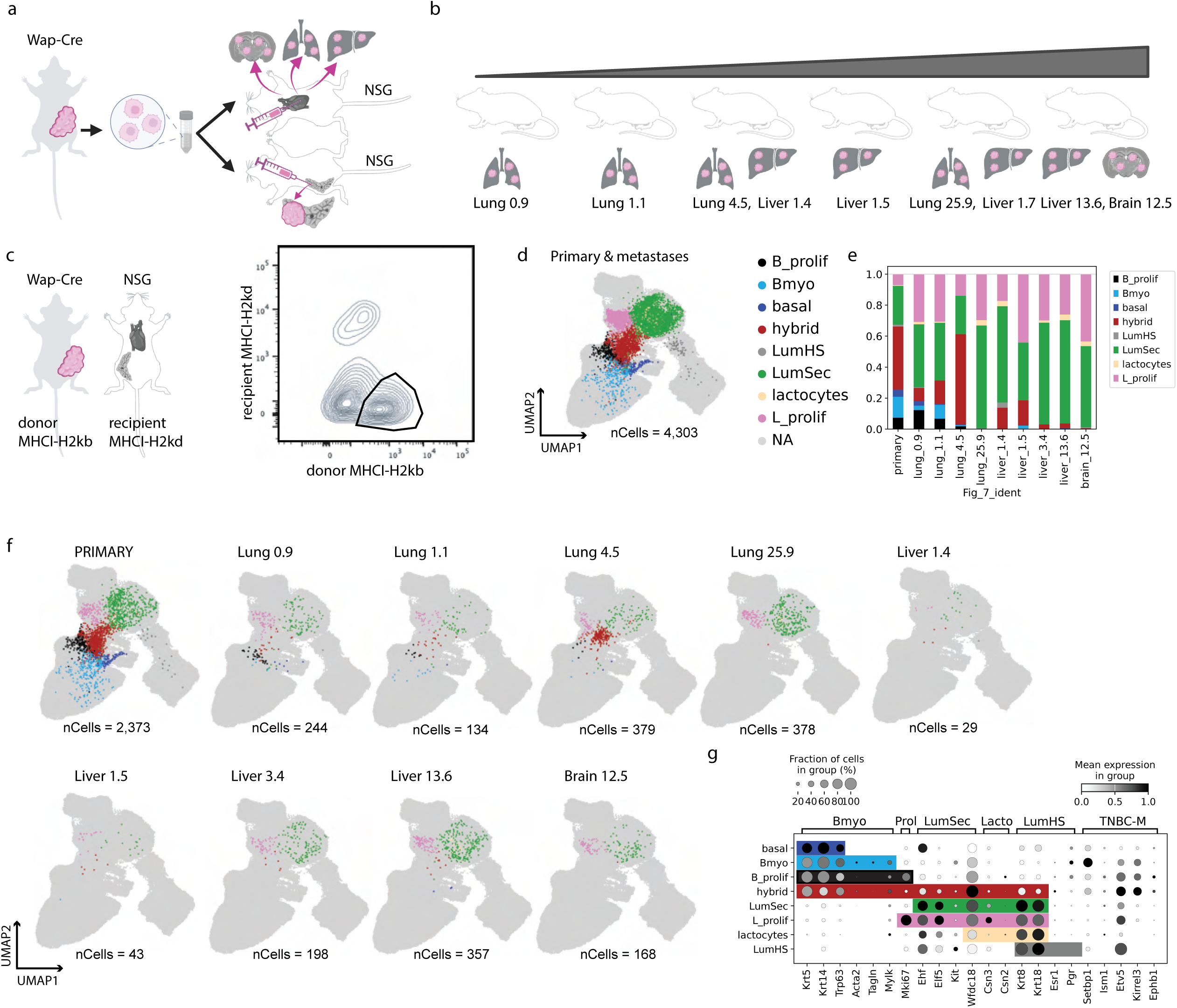
Transcriptomic characterization of cellular states in *Wap-Cre Brca1*-deficient metastatic lesions. **a,** Experimental workflow for experimental metastasis. Primary tumor tissue from a *Wap-Cre* donor was dissociated and intracardially injected into female NSG (NOD-*scid* Il2rg^null^, *n* = 6) recipient mice (*n* = 250,000 cells per injection). **b,** Schematic of resulting metastatic lesions ranked by burden. **c,** Identification and enrichment of metastatic cell fractions using mouse strain-specific antibodies. Left: MHC-I alloantigens (H2-K^b^ vs H2-K^d^). Right: Representative FACS plot showing the isolation of donor tumor cells (H2-K^b+^ H2-K^d-^). **d,** UMAP projection of the joint latent space for the primary tumor and each associated metastatic sample (n = 4,303 cells), colored by Dev-atlas-defined cell populations established in Fig. 5d. **e,** Relative frequencies of Dev-atlas-defined cell populations across the primary and metastatic samples shown in **a,b**. **f,** UMAP projection from **d**, partitioned by primary tumor and each associated metastatic sample. **g,** Dot plots showing expression of canonical MEC markers and derived TNBC-M marker genes (defined in Fig. 2 and **Methods**). Data are stratified by the Dev-atlas-defined cell populations of the metastatic cell fraction. Color intensity represents mean expression level; dot size represents the percentage of cells expressing the gene.

We performed clustering analysis of the primary and metastatic cells using the Dev-atlas integrated embedding. Interestingly, we found that metastases were consistently enriched for luminal-associated states (LumSec, L_prolif) relative to the original primary tumor. We also observed differences in tissues with low versus high burden metastasis. We observed higher frequencies of hybrid cells in lower burden tissues, and higher frequencies of LumSec cells in higher burden tissues (Fig 7d,e). We rarely detected basal cell subsets, and only in low burden lung metastasis samples (Fig. 7d-f and Extended Data Fig. 7c,d). Analysis of canonical marker genes for each epithelial cell subset showed similar expression patterns as observed in primary tumors (Fig. 7g). We also performed single-cell TNBC subtyping analysis of transplanted cells, which showed that the distribution of cells representing the TNBC-M subtype in metastases was indistinguishable from that in the primary and orthotopic tumors (Extended Data Fig. 7e,f), demonstrating that metastatic cells retain gene expression programs associated with TNBC-M despite displaying a shift towards luminal differentiation.

### Cross-species integration identifies human counterparts of the Wap-Cre cell of origin

We investigated how aberrant epithelial differentiation in the *Wap-Cre* and *Blg-Cre* models compares to epithelial differentiation states observed in healthy human breast tissues. We compared differentiation in the *Wap-Cre* and *Blg-Cre* models with a human breast cell atlas dataset (HBCA) containing cells collected from healthy female donors (Fig. 8a)^51^. The HBCA identified one major Bmyo cell population and 7 subpopulations within the LumSec cell type. The 8 HBCA cell states were clearly identifiable after we reprocessed the dataset with our bioinformatic pipeline (Fig 8b). We next performed TNBC-M module scoring on the Bmyo and LumSec cells from the HBCA (Fig. 8b,c). Although scores were low overall, Bmyo cells scored highest for the TNBC-M signature, as in the mouse reference dataset (Fig. 8c). The LumSec-KIT, LumSec-lac, LumSec-myo and LumSec-prol also scored slightly higher for the TNBC-M signature.

**Figure 8.**
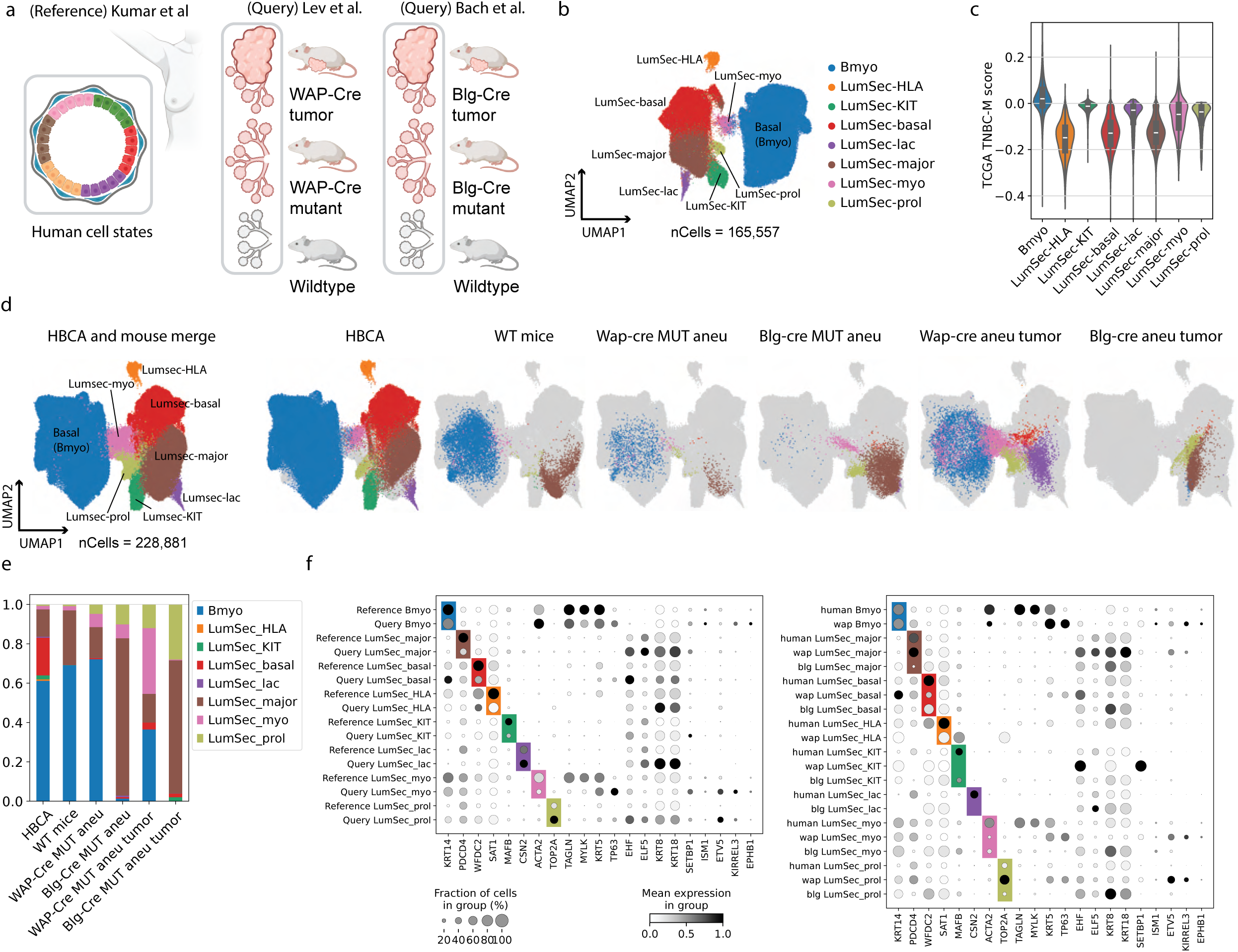
Cross-species integration with the Human Breast Cell Atlas reveals divergent cellular ontogeny of *Brca1*-deficient tumors between mouse models. **a,** Schematic of the experimental design for the transcriptomic integration of the Human Breast Cell Atlas (HBCA) with *Wap-Cre* and *Blg-Cre Brca1*-deficient mouse models. **b,** UMAP projection of the HBCA reference (*n* = 165,557 cells), processed using the computational pipeline described in Fig. 1e and scANVI (see Methods). Cells are colored by original HBCA cell state labels (with “basal” renamed to “Bmyo”). **c,** Violin plots with overlaid box plots displaying human TNBC-M signature enrichment scores across HBCA-defined cell populations. Center line, median; box limits, upper and lower quartiles; whiskers, 1.5x the interquartile range. **d,** UMAP projections of the joint latent space generated by scANVI integration of the HBCA and the *Brca1*-deficient datasets. Cell identities were mapped from HBCA reference using scANVI and partitioned by selected conditions and disease states. **e,** Relative cell-type frequencies within the integrated cross-species dataset, illustrating the distribution of HBCA-defined populations identified through joint scANVI co-clustering across the samples shown in **d**. **f,** Dot plots showing expression of HBCA markers, canonical markers, and derived TNBC-M marker genes (defined in Fig. 2 and Methods). Data are stratified by HBCA-defined cell populations and sub-stratified by experimental batch (left) and selected biological conditions (right). Color intensity represents mean expression level; dot size represents the percentage of cells expressing the gene. Mouse genes were converted to human orthologs for cross-species comparison.

We next integrated the *Wap-Cre* and *Blg-Cre* mouse datasets with the HBCA dataset (Fig. 8a,d). At the level of cell types (Bmyo versus LumSec), integration was highly successful, as determined by the percentage of label agreement before and after integration and co-clustering, both in the mouse datasets and within the HBCA reference itself (Supplementary Fig. 8b-d). All 8 cell states remained identifiable, and the LumSec-myo subset formed an interesting bridge between the basal and luminal cell types. Wild-type mouse cells did not display as much cell state diversity as human cells, with LumSec-major being dominant LumSec cell state and other states barely detectable (Fig. 8d,e). When we compared aneuploid cells from pre-malignant mutant glands (before tumor onset) between the two mouse models, however, the differences were striking: the majority of aneuploid *Wap-Cre* cells clustered with human Bmyo cells, whereas the majority of *Blg-Cre* cells clustered with LumSec-major. Tumors from the two models were likewise dissimilar. While *Blg-Cre* tumor cells were mostly LumSec-major, *Wap-Cre* tumors were composed of the more basal-like cell states. This included Bmyo cells, as well as the two basal-shifted luminal cell states, including LumSec-basal and LumSec-myo, which forms a bridge between basal and luminal cells. As in the mouse developmental integration, *Wap-Cre* cells co-clustering with Bmyo and LumSec-myo did not express myoepithelial genes such as *Acta2*, even though these are defining genes of those states (Fig. 8f). *Wap-Cre* metastases resembled human LumSec cells, co-clustering with LumSec-major, LumSec-myo and LumSec-prol (Extended Data Fig. 8a,b).

We then asked whether human cell states carry different TNBC subtype similarities, and subtyped cell type and cell state pseudo-bulks from the HBCA. The Bmyo pseudo-bulk was classified as TNBC-M-like, whereas the LumSec pseudo-bulk was classified as TNBC-BL1-like (Supplementary Table 7). Within LumSec, individual cell states showed similarity to a variety of TNBC subtypes, and notably the LumSec-myo pseudo-bulk was classified as TNBC-M-like (Extended Data Fig. 8c and Supplementary Table 7). Subtyping of stromal HBCA pseudo-bulks showed that fibroblasts and pericytes also correlate significantly with several TNBC subtypes (Supplementary Table 7), and may therefore contribute to bulk subtype classification and to the discrepancies we observed between tumor-epithelial and tissue-level bulk subtypes in several mouse tumors. Given the similarity of *Wap-Cre* tumors to Bmyo and LumSec-myo cells, and the classification of both of these pseudo-bulks as TNBC-M-like, we propose that mesenchymal subtype tumors originate in these cells.

## Discussion

Basal and luminal cells of origin give rise to distinct TNBC subtypes in *Brca1*-deficient mammary tumors. Previous single-cell analyses of the *Blg-Cre* model identified aberrant luminal differentiation preceding tumor formation^35,36^, a result our reanalysis reproduces: aneuploid cells in mutant *Blg-Cre* glands are LumSec, and *Blg-Cre* tumors are TNBC-BL1. Generalization from the *Blg-Cre* model has established luminal progenitors as the presumed origin of *BRCA1*-associated disease. In *Wap-Cre* mice, Cre recombines both Bmyo and LumSec cells, yet only the basal fraction generates tumors containing basal cells. A second cell of origin therefore exists, and it gives rise to tumors of a different subtype. *Wap-Cre* tumors are basal-derived but mesenchymal, indicating that basal origin does not necessarily specify metaplastic disease and that basal-derived *BRCA1* disease is likely more frequent than the incidence of metaplastic cancer implies. The two models differ not only in the cell populations targeted but also in the Cre driver and in their *Brca1* and *p53* alleles, so the comparison between them does not isolate the cell of origin. Within the *Wap-Cre* model, however, basal and luminal cells from the same donors and carrying identical alleles gave rise to tumors of different phenotype, indicating that lineage contributes independently of genotype.

Somatic mutation is unlikely to specify subtype. Tumor mutational burden is high in *BRCA1*-deficient cancers, yet few recurrent mutations have been identified across large cohorts^6,52,53^. *Wap-Cre* and *Blg-Cre* animals carry loss-of-function *Brca1* alleles yet produce transcriptionally distinct tumors, and *Wap-Cre* tumors converge on the M subtype despite the genomic heterogeneity reported between individual tumors in *Brca1*-driven models^54^. Convergence of this kind is difficult to reconcile with mutational specification of subtype. The mutational landscape of *Wap-Cre* tumors was not profiled here.

The human counterpart of the *Wap-Cre* cell of origin cannot be assigned unambiguously. Epithelial cell states are incompletely conserved between single-cell atlases of the healthy human breast^40,51^, and correspondence between human and mouse states is weaker still. Bmyo and LumSec-myo are both candidates: single-cell TNBC-M-like cells occur in human and mouse Bmyo and LumSec populations, predominantly in Bmyo, and either myoepithelial-related state could lose myoepithelial gene expression to yield the phenotype observed here. Resolution requires lineage tracing in human tissue, which is not currently feasible.

*Wap-Cre* tumors progress from Bmyo cells through a basal state lacking myoepithelial markers to hybrid and LumSec-like states. This trajectory recapitulates fetal mammary differentiation rather than gestation, to which the *Brca1*-mutant phenotype has previously been compared^35,37^. Pre-malignant glands contain predominantly Bmyo and LumSec cells and few hybrid or basal cells, indicating de-differentiation of Bmyo cells rather than expansion of a residual fetal population. Whether the fetal-like state is permissive for transformation or a consequence of it is unresolved, and will require lineage tracing from the Bmyo compartment through tumor onset.

These conclusions carry several limitations. Single-cell genotyping requires physical material and could not be applied to the published *Blg-Cre* dataset; the conclusion that only LumSec cells transform in that model rests on copy-number inference. The CODEX panel does not distinguish developmental basal cells from Bmyo cells that have lost myoepithelial markers, localizing the aberrant population without resolving its identity. The metastasis experiment derives from a single donor tumor in immunodeficient recipients, and intracardiac injection bypasses the early steps of the metastatic cascade; it therefore reports differentiation state rather than metastatic competence.

An M-like transcriptional program was retained across tumor size, differentiation state and tissue environment, persisting through progression, transplantation and metastasis, consistent with specification at initiation. Bulk classification is nonetheless sensitive to stromal composition: fibroblast and pericyte pseudo-bulk profiles correlate with TNBC subtypes and can drive tissue-level assignments that diverge from those of the tumor epithelium. Single-cell subtyping is not subject to this confounder and tracked the cell of origin in both models. Baseline single-cell subtype distributions varied between models and between species, and tumors are therefore best evaluated against the baseline of their own dataset.

Subtype stability is not universally reported. In matched primary and metastatic TNBC, most patients retained their molecular subtype, but mesenchymal tumors frequently transitioned to another subtype at recurrence or metastasis^32^. Several factors could account for the discordance. The stromal contribution to bulk assignment described above could itself differ between primary and metastatic sites. The metastasis experiment reported here derives from a single primary tumor and involved no therapeutic intervention, and therefore samples neither inter-tumor variation nor treatment-associated selection.

Cell of origin carries implications beyond classification. Disease in *BRCA1* carriers can initiate in more than one epithelial compartment: LumSec cells in the *Blg-Cre* model and Bmyo cells in the *Wap-Cre* model. A prevention strategy directed at luminal progenitors alone would therefore leave the basal compartment unaddressed, and effective prevention will need to encompass every epithelial cell type with transformation potential.

## Methods

### Mouse Models and Ethics Statement

All animal experiments were conducted in accordance with the University of California, Irvine, Institutional Animal Care and Use Committee (IACUC) protocol AUP-22-033. Mice were housed in a specific pathogen-free (SPF) facility under a 12-hour light/dark cycle with free access to food and water.

The *Wap-Cre Brca1^f10/f10^p53^f5&6/f5&6^*strain (originally provided by Eva Y.-H. P. Lee) was maintained on a mixed 129/SvJ;C57BL/6 genetic background. Exon numbering for the floxed *Brca1* alleles follows the current GRCm38 (mm10)/GRCm39 (mm39) annotation, under which the *Wap-Cre* allele targets *Brca1* exon 10 and the *Blg-Cre* allele targets *Brca1* exons 21–23. The original reports of these alleles designated them exon 11 and exons 22–24, respectively, under the genome builds available at the time; the targeted sequences are unchanged. Female littermates or age-matched, co-housed females were utilized for all experimental groups to minimize environmental variance. For physiological and transcriptomic analysis, “wild-type” and “mutant” cohorts were analyzed at 4 months of age; “tumor” cohorts were analyzed between 4 and 6 months of age. Transplantation experiments utilized female NOD *scid* gamma (NSG, JAX 005557) mice as recipients. Recipient ages were restricted to 4–5 weeks (prepubescent) for mammary fat pad transplants and 6 weeks for experimental metastasis assays. Only female mice were used in this study due to the sex-specific prevalence of breast cancer and the mammary-directed expression of the *Wap-Cre* transgene.

### Genotyping

Genomic DNA was extracted from tail biopsies at weaning. PCR amplification was performed using the following primers:

***Trp53* Forward:** 5’-TGG GAC AGC CAA GTC TGT TA-3’

***Trp53* Reverse:** 5’-CAT GCA GGA GCT ATT ACA CA-3’

***Cre* Forward:** 5’-GGT GTC CAA TTT ACT GAC CGT ACA-3’

***Cre* Reverse:** 5’-CGG ATC CGC CGC ATA ACC AGT G-3’

Amplification of the Trp53 locus yielded an approximately 500 bp product for the floxed Trp53 allele, and a 363 bp product for the Cre transgene.

### Tissue Harvest and Dissociation

Mice were anesthetized with isoflurane, and lymph nodes were labeled with Evans blue dye to aid in removal. Animals were euthanized via CO_2_ inhalation followed by cervical dislocation. To remove peripheral blood, mice underwent cardiac perfusion with PBS-EDTA buffer. After lymph node removal, mammary fat pads (MFPs) 1–5 were harvested; contralateral glands were pooled to yield five distinct MFP samples per animal. In mice with palpable tumors, the tumor was harvested and the remaining MFP tissue was discarded.

Tissues were mechanically minced into approximately 1 mm^3^ fragments using a sterile razor blade and enzymatically digested with Collagenase IV (2 μg/mL, Sigma-Aldrich cat. no. C5138-1G) in DMEM (Corning, 10-013-CV), followed by treatment with DNase I (Worthington Biochemical Corporation, cat. no. LS002139) and 0.25% Trypsin-EDTA (Corning, 25-052-Cl), with intervening chilled PBS washes. Resulting single-cell suspensions were filtered through a 70 μm strainer, cryopreserved in [45% DMEM, 45% FBS, 10% DMSO], and batched for subsequent analysis.

### Mammary Fat Pad Transplantation

Donor cells from 4-month-old *Wap-Cre Brca1^f10/f10^p53^f5&6/f5&6^* mice were isolated and prepared as described above. For FACS-based isolation (FACSAria Fusion, BD Biosciences), cells were stained with antibodies against CD45, EpCAM, and CD49f. Cells were gated as follows: basal (Cd45^neg^ EpCAM^med^ Cd49f^high^), luminal (Cd45^neg^ EpCAM^high^ CD49f^med^), and stromal (CD45^neg^ EpCAM^neg^ CD49f^neg^) after doublet and debris exclusion.

For transplantation, 12,500 sorted epithelial cells were mixed with 12,500 stromal cells and resuspended in a 1:1 (v:v) mixture of PBS and Growth Factor Reduced Matrigel (Corning 356230) for a total of 25,000 cells in 20 μL. Recipient 4–5 week-old female NSG mice underwent bilateral clearing of the #4 MFP under isoflurane anesthesia, as previously described^55^. Carprofen (5 mg/kg) was administered intraperitoneally for analgesia. Resulting tumors were collected when the largest tumor reached the endpoint size of 25mm in diameter. Successful engraftment was confirmed by the presence of epithelial outgrowths in non-tumorous glands. Two independent experimental replicates were performed using distinct donor pools.

### Experimental Metastasis and Enrichment

A late-stage endpoint tumor (Sample ID tumor_5_25mm) was utilized for both metastasis assays, which were performed in separate cohorts of recipient mice. For experimental metastasis, 250,000 cells in 100 μL PBS were injected into the left ventricle of each recipient (n = 6) via ultrasound guidance (Butterfly iQ ultrasound device, Butterfly Network, Inc.). A separate control cohort (n = 2) received the same number of cells orthotopically into the left #4 MFP. No animal received both injections. Anesthesia and analgesia were administered as described for the MFP transplantation assay.

Tissues were collected after 4 weeks. Single-cell suspensions from lungs, livers, and orthotopic tumors were prepared and cryopreserved as described in the tissue harvest and dissociation section. Dissected brains were partitioned and transferred into C Tubes (Miltenyi Biotec, cat. no. 130-093-237) containing HBSS (with Ca^2+^ and Mg^2+^) supplemented with 1 mg/mL Collagenase D (Millipore Sigma, cat. no. 11213857001) and 10 μg/mL DNase I (Fisher Scientific, cat. no. NC9709009). Tissues were dissociated using the gentleMACS Octo Dissociator with Heaters following the manufacturer’s 37°C adult brain dissociation protocol (ABDK1_37C). Digested suspension was filtered through a 70 μm cell strainer. Myelin was removed via density gradient centrifugation using Debris Removal Solution (Miltenyi Biotec, cat. no. 130-109-398) according to the manufacturer’s instructions. Residual red blood cells were eliminated using ACK Lysis Buffer (Gibco, cat. no. A10492-01) for 2 minutes on ice. The final single-cell suspension was washed in chilled FACS buffer and cryopreserved in [45% DMEM, 45% FBS, 10% DMSO].

To identify samples suitable for single-cell analysis, metastatic burden was first screened via flow cytometry. Single-cell suspensions were stained for donor-specific H-2K^b^ (PE-conjugated, AF6-88.5.5.3, Invitrogen cat. no. 12-5958-82), and host-specific H-2K^d^ (APC-conjugated, SF1-1.1.1, Invitrogen cat. no. 17-5957-82) MHC markers. Metastatic burden was quantified as the frequency of donor-derived cells H-2K^b+^ H-2K^d-^ expressed as a percentage of total viable, non-doublet cells. Only samples with a detectable metastatic burden >0.5% were selected for subsequent enrichment and sequencing.

For scRNA-seq, metastatic cells were enriched via positive selection using the EasySep PE Positive Selection Kit II (STEMCELL Technologies cat. no. 17684). Briefly, cells were labeled with the aforementioned H-2K^b^ and H-2K^d^ antibodies and subjected to three sequential rounds of immunomagnetic positive selection. Following enrichment, cells were washed and multiplexed using 10x Genomics CellPlex CMO tags (User Guide CG000391). To ensure maximum purity and viability, the enriched fraction was further refined by FACS (FACSAria, BD Biosciences) to isolate viable (Sytox Blue negative) single cells and proceeded immediately to library construction. As a control for the effect of immunomagnetic selection on recovered cell composition, an aliquot of the donor primary tumor (tumor_5_25mm) was carried through the identical enrichment, multiplexing and sorting workflow and sequenced alongside the metastatic samples, yielding 757 cells; the same tumor had also been sequenced without enrichment as part of the primary dataset (3,147 cells; Fig. 1e). Enriched and unenriched profiles were indistinguishable and were combined for downstream analysis, contributing 757 and 1,616 epithelial cells, respectively, to the metastasis analysis (Fig. 7). The metastasis experiment yielded 4,806 cells after quality control, bringing the total number of cells sequenced in this study to 67,001.

### Single-cell RNA-seq Library Preparation and Multiplexing

Cryopreserved single-cell suspensions were thawed rapidly at 37 °C, filtered through a 70 μm cell strainer, and washed twice in chilled PBS. Cell concentration and viability were assessed using a Countess II Automated Cell Counter (Thermo Fisher Scientific). To enable sample multiplexing, up to 2 x 10^6^ cells per sample were labeled with 10x Genomics CellPlex Cell Multiplexing Oligos (CMOs) according to the manufacturer’s instructions (Demonstrated Protocol CG000391). Following labeling, samples were pooled and subjected to FACS-based debris removal and live-cell enrichment (Sytox Blue negative). To ensure balanced representation across multiplexed libraries, similar cell numbers from each CMO-labeled sample were sorted into a single collection tube containing chilled FACS buffer (PBS supplemented with 10% FBS). The final cell concentration and viability were re-verified before proceeding to 10x Chromium GEM generation.

### scRNA-seq Library Construction and Sequencing

Single-cell suspensions were loaded onto the Chromium Controller X (10x Genomics) using the Chromium Next GEM Single Cell 3’ Kit (v3.1 chemistry). We targeted a capture of up to 30,000 cells per lane (approximately 2,500 cells per CMO tag). cDNA amplification and library construction (including Gene Expression and Cell Multiplexing libraries) were performed according to the 10x Genomics User Guide (CG000388, Rev C) without modification. Final library quality and fragment size distributions were assessed using the Agilent Bioanalyzer High Sensitivity DNA kit. Libraries were pooled and sequenced on an Illumina NovaSeq 6000 by the UC Irvine Genomics and Technology Hub, aiming for a depth of at least 15,000 read pairs per cell.

### Computational Processing and Demultiplexing

Raw sequencing data (BCL files) were demultiplexed using bcl2fastq (v2.20, Illumina) by the UC Irvine Institute for Genomics and Bioinformatics. Subsequent processing was performed using Cell Ranger (v7.0.1, 10x Genomics)^56^. Reads were aligned to the mm10 reference genome (version 2020-A). Sample demultiplexing and CellPlex Oligo (CMO) assignment were executed using the cellranger multi pipeline to identify individual cell barcodes and assign sample identities.

Downstream analysis was conducted using a combination of Scanpy (v1.10.1) in Python^57^ and Seurat (v5.0.3) in R^58^, with metadata and count matrices exchanged via .h5ad and .rds formats to leverage specialized tools in both ecosystems.

### Data Quality Control and Ambient RNA Removal

Initial quality control and filtering were performed independently for each sequencing batch. Ambient RNA contamination was corrected using SoupX (v1.6.2), with contamination fractions adjusted in integer increments to preserve the integer-based count matrix required for downstream feature selection^59,60^. To account for batch-specific technical variation, we applied cluster-specific adaptive thresholds for cell filtering; cells with aberrant total counts (*n*UMI) or feature counts (*n*Gene) were identified by inspecting the tails of the distribution within each cluster and excluded. Mitochondrial read thresholds were set at 5–8% depending on the specific batch distribution.

### Doublet Detection and Consensus Filtering

A multi-step consensus strategy was employed to ensure high-fidelity single-cell resolution. First, inter-sample multiplets were identified and excluded during the cellranger multi demultiplexing step based on the presence of multiple CellPlex Cell Multiplexing Oligos (CMOs). To address intra-sample doublets, we utilized DoubletFinder (v2.0.3)^61^ to calculate doublet scores for each batch independently. While the expected doublet rate was considered, we prioritized a biological consensus for final exclusion. Following batch merging, formal doublet sub-clusters were removed only when they met three rigorous criteria: (1) high DoubletFinder scores, (2) significantly elevated *n*UMI/*n*Gene counts, and (3) co-expression of mutually exclusive canonical markers (e.g., *Epcam* and *Ptprc*). For batch #1, which was loaded at a higher cell density, an initial round of doublet exclusion was performed prior to integration.

### Dimensionality Reduction and Clustering

High-quality cells were processed through a standardized pipeline. First, non-expressed genes were removed, and highly variable genes (HVGs) were identified using the BigSur algorithm^59,60^ within the Scanpy framework to ensure robust feature selection. Data were normalized to total counts per cell, and sequencing depth was regressed out prior to scaling. Principal Component Analysis (PCA) was performed on the HVG set, and the first 40 PCs were used to construct a nearest-neighbor graph. Clustering was performed using the Leiden algorithm^62^. A resolution of 0.1 was applied for initial individual batch assessment, while a resolution of 1.0 was used for the merged dataset to resolve fine-grained cell states. Uniform Manifold Approximation and Projection (UMAP)^63^ was utilized for two-dimensional visualization. For all sub-clustering analyses (e.g., epithelial-specific refinement), the entire pipeline from HVG selection to embedding was re-executed to capture compartment-specific variance.

### Integration and Batch Correction

To mitigate technical batch effects while preserving biological variation, we employed scVI (scvi-tools v1.1.2)^64,65^ for data integration. Batch correction was applied specifically during sub-clustering of the epithelial compartment, where inter-tumor heterogeneity otherwise drove batch-specific clustering. The resulting latent space was used for downstream UMAP visualization and clustering.

### Cell-Type Annotation and Quality Control Refinement

Initial cluster identities were manually assigned by querying the expression of canonical lineage-specific marker genes (same as shown in Extended Data Figure 1c). Following initial annotation, a strict post-sequencing quality control filter was applied: two wild-type mammary fat pad (MFP) samples out of fifteen were bioinformatically excluded from all downstream analyses due to an anomalous, extreme enrichment of B-cells and T-cells, indicating incomplete excision of adjacent lymph nodes during tissue harvest. For all subsequent re-clustering and compartment-specific sub-clustering (e.g., epithelial lineage isolation), cells were subsetted, and the entire computational workflow—including highly variable gene selection, data normalization, dimensional reduction, and graph-based clustering was re-executed.

### Inference of Copy-Number Variations and Clonal Substructure

Chromosomal copy number variations (CNVs) and aneuploidy profiles were inferred from single-cell transcriptomes using the CopyKAT (v1.1.0) algorithmic framework^41^, adapted for mouse data by setting the genome configuration to mm10. To manage computational density while preserving statistical power, only epithelial datasets (excluding the Luminal-HS population) were used. Highly purified epithelial cells isolated from two distinct wild-type mice served as baseline normal references, while a third chronological mouse and a pooled normal epithelial control population obtained via fluorescence-activated cell sorting (FACS) were utilized as non-malignant controls.

To mitigate technical batch artifacts, to which single-cell CNV inference is highly sensitive, cells from the batch #1 tumor (which experienced GEM-generation overloading and subsequent low sequencing coverage) were isolated and analyzed independently; for this specific run, intra-sample stromal cells were leveraged as the internal, known diploid reference baseline.

### Differential Abundance Analysis via Cellular Neighborhoods

To identify continuous cellular states undergoing significant frequency shifts between experimental conditions without the bias of discrete cluster boundaries, we performed differential abundance analysis using the Milo^38^ (v1.4.0) R package. Single-cell object formats were converted from Seurat to SingleCellExperiment structures prior to construction of k-nearest neighbor (KNN) graphs. Neighborhood definitions and graph topologies were constructed using the following specific parameterizations:

The graph was built using the top 20 principal components (instead of the default 30) for the buildGraph, makeNhoods, and calcNhoodDistance functions.

The neighborhood sampling proportion (prop) was set to 0.25 within makeNhoods to capture fine-grained topological density.

All remaining parameters were maintained at package defaults. Differential abundance testing across the resulting cellular neighborhoods was modeled using generalized linear models within edgeR^38,66^, and statistical significance was determined via a Quasi-Likelihood *F*-test (testNhoods). Neighborhood-level log-fold changes and false discovery rates (FDR) were visualized via beeswarm projections (plotDAbeeswarm).

### Bimodal Gene Expression Scoring along the Basal–Luminal Axis

To quantify the phenotypic position of epithelial cells along the differentiation hierarchy, a continuous differentiation axis score was developed. First, highly resolved lineage-specific marker gene lists were established for wild-type basal-myoepithelial (Bmyo) cells and luminal-secretory (LumSec) cells using Scanpy’s rank_genes_groups function; genes were retained only if they exhibited a log-fold change ≥ 1.5 and a Benjamini-Hochberg adjusted *P*-value < 0.05. Gene lists were generated from wild-type *Wap*-driven epithelial cells and are provided in Supplementary Table 3.

Module scores for the top 500 lineage-specific marker genes were calculated across the epithelial subset (excluding the Luminal-HS cluster) using Scanpy’s score_genes function. To ensure cross-species comparability, any marker gene absent from any of the target evaluation datasets was uniformly omitted from all scoring matrices, resulting in 471 LumSec and 480 Bmyo genes (Supplementary Table 3). The resulting raw module scores were mathematically scaled to a continuous **[-1, +1]** interval. The basal score vector was negated, and the final consolidated axis score ***S*_axis_**was derived by calculating the sum of the luminal score ***S*_luminal_**and the negated basal score **-*S*_basal_,** followed by a final linear rescaling to a uniform **[-1, +1]** scale:

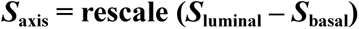

where a final value of -1.0 represents a pure basal-myoepithelial state and +1.0 represents a fully committed luminal-secretory phenotype.

### Generation of Transcriptomic Tumor Signatures

Lineage-specific malignant activation signatures for the *Wap-Cre* and *Blg-Cre* models were constructed using Scanpy’s rank_genes_groups tool by comparing tumor cells against non-malignant wild-type epithelial controls. To establish an unbiased reference baseline, wild-type epithelial profiles were randomly downsampled to 3,000 cells maintaining a strict 1:1:1 ratio of Basal : Luminal-Secretory : Luminal-HS states. The *Blg-Cre* tumor matrix comprised all 3,052 recovered tumor cells. To avoid statistical power imbalances, the *Wap-Cre* tumor compartment was selectively downsampled to match this exact depth (n = 3,052 cells total); individual *Wap-Cre* tumors possessing fewer than 690 cells were included in their entirety, while larger tumors were randomly downsampled to exactly 690 cells per sample. Only mesenchymal *Wap-Cre* tumors were utilized. Differentially expressed genes were filtered to retain robust candidates exhibiting a Log_2_ fold change ≥ 1.5 and an adjusted *P*-value < 0.05.

Mesenchymal (TNBC-M) transcriptomic tumor signature was similarly derived using primary tumor RNA-sequencing matrices from The Cancer Genome Atlas (TCGA) BRCA cohort. To isolate a rigorous clinical TNBC population, analysis was restricted to samples possessing a Basal-like subtype designation that successfully passed the *ESR1* expression thresholding filter within the Vanderbilt TNBCtype stratification framework^43^. Expression count matrices were converted into Scanpy AnnData objects for downstream differential expression testing. To define the Mesenchymal (M) lineage signature, Scanpy’s rank_genes_groups function was executed to contrast M-subtype tumors against all non-mesenchymal TNBC profiles. Candidate differentially expressed genes were filtered using identical stringent parameters to those described above, retaining targets with a Log_2_ fold change ≥ 1.5 and an adjusted *P*-value < 0.05.

### Triple-Negative Breast Cancer Subtyping and Homology Mapping

Bulk and pseudo-bulked transcriptomic profiles were stratified into molecular subtypes using the web-based TNBCtype classifier (https://cbc.app.vumc.org/tnbc/)^43^. Human bulk RNA-seq count matrices from triple-negative breast cancer (TNBC) patients were log-transformed and normalized via the trimmed mean of M-values (TMM) method. Mouse single-cell profiles were aggregated into biological pseudo-bulk matrices (summing counts across all cell types per individual mouse) and similarly TMM-normalized. Mouse genes were mapped to their corresponding human orthologs using the Mouse Genome Informatics (MGI) Homology Suite^67^. Samples classified into highly heterogeneous or non-epithelial categories—specifically Immunomodulatory (IM), Mesenchymal Stem-Like (MSL), or Unclassified (UNS)—were manually assigned to their next-highest positive-scoring primary epithelial centroid: Basal-like 1 (BL1), Basal-like 2 (BL2), Luminal Androgen Receptor (LAR), or Mesenchymal (M). Samples presenting strictly negative centroid correlations across all target lineages were designated as Unclassified Epithelial (UNC) as described previously^28^. Clinical tumors that were pathologically confirmed as TNBC but initially rejected by the automated classifier due to residual *ESR1* baseline transcripts were re-analyzed and subtyped following computational filtering of the *ESR1* expression values.

### Transcriptomic Scoring of Clinical TNBC Cohorts

To characterize how our murine tumor signatures align with the clinical spectrum of human triple-negative breast cancer (TNBC), we analyzed two independent human clinical cohorts. The first cohort was a multi-institutional dataset obtained from the Basser-Gray Foundation Repository (BGFR), which was curated and shared by the Nathanson Laboratory at the University of Pennsylvania. Of these, 17 samples were previously described^24^ and 34 are newly reported here. The second cohort was the breast carcinoma cohort from The Cancer Genome Atlas (TCGA). For the BGFR cohort, human *BRCA1*-mutated and sporadic TNBC bulk RNA-seq expression profiles and linked clinical annotations were compiled into a Seurat (v5) framework and normalized using the NormalizeData function with default configurations. For the TCGA cohort, normalized TCGA breast invasive carcinoma (TCGA-BRCA) RNA-seq data (Log_2_-transformed RSEM normalized expression values) and corresponding clinical metadata were downloaded via the UCSC Xena Browser. Samples across both datasets were subsequently filtered to isolate the pathologically and transcriptomically confirmed TNBC subpopulation to analyze human subtype distributions.

Mouse-derived *Wap-Cre* and *Blg-Cre* tumor signatures were mapped to their corresponding human orthologs using the Mouse Genome Informatics (MGI) Homology Suite. Human patient transcriptomes within the BGFR cohort were scored against these humanized signatures using Seurat’s AddModuleScore function, computing expression metrics relative to a control feature set composed of 100 randomly selected genes. To evaluate signature distributions across clinical TNBC variants, patients within each cohort were partitioned based on their TNBCtype classification: Mesenchymal (TNBC-M) versus non-mesenchymal lineages (TNBC-Other: BL1, BL2, LAR, UNC).

Following score computation, the clinical expression matrices and calculated module score metadata from both datasets were exported and converted into Scanpy AnnData objects to facilitate Python-based statistical testing and plotting. Statistical differences in mean module scores between these two primary groups were evaluated using a one-way analysis of variance (ANOVA) followed by Tukey’s post-hoc test utilizing pingouin and scipy.stats in Python^68,69^. Final visualizations of the clinical module scores stratified by subtype were generated using matplotlib and seaborn to maintain graphic and aesthetic consistency across all manuscript figures.

### Single-Cell Target Genotyping

Single-cell genotyping of endogenous somatic alterations was performed by adapting the Mutation Enrichment via RNA single-cell sequencing (MAESTER) framework^44^. Due to the large genomic span of the *Brca1* locus preventing efficient multiplexed PCR amplification, transcript expression of *Trp53* was leveraged as a proxy, as both alleles are regulated downstream of the shared mammary-specific *Wap* promoter.

Targeted amplification of *Trp53* transcripts was executed using 20 ng of 3′-barcoded, full-length cDNA libraries generated via standard scRNA-seq. PCR1 was performed using a locus-specific *Trp53* forward primer:

5′-CACCCGAGAATTCCATGGGACAGCCAAGTCTGTTA-3′

paired with a barcoded GoT-P5-i5-BCXX reverse primer to append sample-specific indexing configurations. PCR1 cycling parameters were modified to optimize target fidelity, decreasing the annealing temperature to 60°C and extending the annealing duration to 90 seconds.

PCR2 was subsequently performed to attach full-length Illumina sequencing adaptors and dual 10-base-pair library indexes for multiplexing; the PCR2 annealing step was extended to 5 minutes. Amplified fragments were purified via solid-phase reversible immobilization using SPRIselect beads (Beckman Coulter). Final enriched genotyping libraries were multiplexed and sequenced on an Illumina MiSeq Micro flow cell, targeting 4 million reads in a 300-bp paired-end (300PE) configuration.

### Reference Atlas Curation and Cross-Species Integration

#### Mouse Mammary Reference Construction and Unified Training

To project query *Brca1*-mutant mammary tumor datasets into a baseline, non-malignant physiological context, a public single-cell mouse mammary gland reference dataset^45^ was obtained from the Gene Expression Omnibus (GEO: GSE164017). Background mouse strains irrelevant to our experimental models were bioinformatically excluded, restricting the baseline reference atlas strictly to C57BL/6 and FVB genetic backgrounds. Reference data quality control was refined via visual inspection of baseline distributions: stromal cell contaminants were discarded, and low-quality cells were removed if they exhibited greater than 10% mitochondrial transcript content, greater than 6,000 unique detected genes, or greater than 50,000 total reads.

To ensure absolute cell-type annotation consistency across our datasets, the filtered mouse reference atlas was processed and manually annotated de novo using the exact same computational pipeline, marker genes, and cluster-curation criteria established for our experimental *Wap-Cre* data analysis. To prepare this re-annotated mouse reference atlas for supervised cell-type mapping, an operational variational framework was constructed using the scvi-tools (v1.1.2) suite. Initial batch correction across distinct source datasets was performed by training a variational autoencoder (scVI) using the orig.ident metadata slot as the categorical batch covariate, restricting feature selection to the top 838 highly variable genes (HVGs) identified by the BigSur algorithm^59,60^. Following latent representation modeling, a conditional variational autoencoder (scANVI)^65,70^ was constructed, leveraging our newly assigned lineage definitions as reference labels to establish a fully trained cell-type classification model.

#### Human Breast Cell Atlas Co-Embedding and Mapping

To cross-evaluate the translational cellular states of our murine models against normal human tissue architecture, single-cell transcriptomic reference profiles from the Human Breast Cell Atlas (HBCA^51^; GEO: GSE235326) were incorporated into the reference mapping pipeline. Unlike the mouse atlas, the human dataset was not re-annotated; instead, the original cell-type labels defined by the authors were strictly retained. To establish a unified feature space, mouse gene symbols from our experimental datasets and the custom-annotated mouse reference atlas were mapped to their corresponding human orthologs using the Mouse Genome Informatics (MGI) Homology Suite.

Epithelial single-cell matrices from the experimental *Wap-Cre* and *Blg-Cre* cohorts were designated as query datasets and restricted to the same highly variable human orthologs defined by the reference model architecture; notably, the *Luminal-HS* cell population was bioinformatically excluded from this cross-species analysis. To achieve cross-species alignment, reference profiles (both the human baseline HBCA dataset and the custom-annotated, humanized murine reference) and our experimental tumor query matrices were integrated via latent space concatenation utilizing the trained scANVI conditional autoencoder.

The resulting integrated latent spaces—modeled independently for the mouse and human reference tracks—were co-embedded using Uniform Manifold Approximation and Projection (UMAP) and co-clustered via the Leiden algorithm. This strategy of evaluating shared co-cluster coordinates served as the definitive method for annotating query cells within both reference frameworks. Final query cell-type annotations for both the mouse and human mappings were assigned based on the dominant reference label within each shared co-cluster coordinate. While direct scANVI automated label-transfer prediction probabilities were continuously monitored in both pipelines to verify classification confidence, these discrete, co-clustering consensus labels were strictly utilized for all downstream analytical workflows, cell-type assignments, and cross-species state comparisons.

#### Co-Detection by Indexing (CODEX) Spatial Profiling

Formalin-fixed paraffin-embedded (FFPE) mammary gland and tumor tissues from *Brca1*-wildtype and mutant cohorts were analyzed using the PhenoCycler (CODEX) platform (Akoya Biosciences). Tissues were sectioned at 5 μm thickness, mounted onto poly-L-lysine-coated glass coverslips, and processed per the manufacturer’s protocol. A multiplexed panel of antibodies (Supplementary Table 6) was utilized for staining, with targets and oligo-barcode conjugations specified in the supplementary materials. Iterative imaging cycles were executed on the PhenoCycler-Open system, with fluorescent signal acquisition performed on a Keyence BZ-X800 fluorescence microscope equipped with a 20X PlanApo (0.75 NA) objective. Raw image data were processed and stitched using the CODEX Processor (v1.8).

#### Spatial Segmentation and Single-Cell Feature Quantification

Tissue segmentation was performed using the StarDist deep-learning framework within QuPath^71,72^ to generate high-fidelity nuclear masks anchored by DAPI signal intensity. To ensure architectural reproducibility, cell detection was executed using the standard, pre-trained StarDist 2D model (dsb3_2018) without additional fine-tuning. Protein expression profiles were quantified by calculating the mean intensity within defined cellular compartments: nuclear-localized proteins (including Ki-67, P63, and PR) were quantified within the primary StarDist nuclear mask, whereas membrane- or cytoplasmic-localized targets (including SMA, K8, K14, and K5) were quantified using an expanded membrane mask.

#### Computational Analysis of Spatial Proteomic Data

Due to the highly eccentric, elongated cytoplasmic morphology of basal-myoepithelial (Bmyo) cells making robust whole-cell segmentation technically challenging, downstream spatial analyses were strictly restricted to nuclear compartment coordinates to eliminate cytoplasmic boundary overlapping artifacts. Segmented single-nucleus feature matrices containing raw marker expression levels and exact Cartesian (X, Y) spatial coordinates were loaded into Seurat (v5) utilizing the ReadAkoya pipeline.

To prevent technical batch confounding and preserve sample-specific spatial architectures, the standard Seurat spatial analysis vignette for Akoya data was executed independently for each of the nine biological replicates—comprising three wild-type, three mutant, and three tumor tissue samples.

All average protein intensity values were z-score normalized across the single-nucleus population of each sample independently. Unsupervised cell-state classification was conducted via the Leiden algorithm with the resolution parameter set between 0.25 and 0.5 depending on sample complexity, and cluster labels were assigned to cells within each sample independently.

Following independent sample processing, quality control filtering, and annotation, the individual Seurat objects were merged into a unified spatial matrix. To ensure structural and stylistic consistency across all manuscript visualizations, the combined single-nucleus expression matrix, along with linked metadata and coordinate slots, was converted into a Scanpy AnnData object structure. All final multi-replicate marker intensity distributions were generated within the Scanpy framework to match the formatting of the single-cell transcriptomic figures.

#### Standard Immunofluorescence

Standard immunofluorescence was performed on 5μm FFPE sections. Sections were deparaffinized in Histo-Clear (National Diagnostics) and rehydrated through a graded ethanol series. Antigen retrieval was achieved via microwave-based heat-induced epitope retrieval (HIER) in 10 mM citric acid buffer (pH 6.0, 0.05% Tween-20). Tissues were blocked using BlockAid (Thermo Fisher Scientific), incubated with primary antibodies overnight at 4°C, and subsequently probed with species-specific secondary antibodies for 1 hour at room temperature. Slides were mounted in VECTASHIELD containing DAPI (Vector Laboratories). Fluorescent image acquisition was conducted on a Keyence BZ-X700 microscope.

### Statistics and Reproducibility

Bar plots show the mean ± s.e.m. across the biological replicates defined in the corresponding figure legend. For box plots, the center line represents the median, box limits the upper and lower quartiles, and whiskers 1.5× the interquartile range. Differences in module scores between clinical TNBC subtype groups were evaluated by one-way analysis of variance (ANOVA) followed by Tukey’s post-hoc test, implemented in pingouin and scipy.stats. Differences in tumor-formation frequency between transplanted basal and luminal fractions were evaluated by a two-sided Fisher’s exact test. This test treats injection sites as independent observations. Eight recipients received contralateral basal and luminal implants and one received bilateral basal implants, so sites within a recipient may not be fully independent. The comparison is therefore reported as a site-level analysis, and residual within-mouse dependence is a limitation of this estimate. All calculated numerical *P* values and statistical test specifications are indicated directly within the corresponding plots and legends. Transplantation and imaging experiments were performed with at least two independent biological replicates, as specified in the corresponding figure legends. No statistical frameworks or power analyses were deployed to pre-determine sample sizes; sample sizes were chosen based on historical empirical benchmarks for single-cell transcriptomic and murine transplantation assays.

Age-matched female NSG mice were bred in-house and randomly allocated into experimental groups upon reaching the required chronological age. Blinding was not enforced; investigators were aware of the distinct genetic backgrounds and experimental conditions during animal handling, tissue harvesting, and downstream computational data analysis.

### Data Availability

The raw and processed single-cell RNA-sequencing (scRNA-seq) data generated in this study have been deposited in the National Center for Biotechnology Information (NCBI) Gene Expression Omnibus (GEO) and are accessible under accession number GSE338147.

The clinical human triple-negative breast cancer (TNBC) bulk RNA-seq datasets analyzed during this study were obtained from the Basser-Gray Foundation Repository (through the Nathanson Laboratory at the University of Pennsylvania); restrictions apply to the public availability of these clinical datasets, which were used under license for the current study, but are available from the corresponding author upon reasonable request. The *Blg-Cre* model data utilized for comparison is available at http://marionilab.cruk.cam.ac.uk/BRCA1Tumourigenesis. The public mouse mammary gland single-cell reference dataset utilized for atlas mapping is available via GEO under series accession GSE164017.

Raw and processed multiplexed imaging datasets, including stitched image tiles, StarDist segmentation masks, and single-nucleus feature matrices generated via Co-Detection by Indexing (CODEX), and source data for Figure 6 are available from the corresponding author upon reasonable request.

### Code Availability

Analysis code is available in the project’s GitHub repository: https://github.com/Dlawsonlab/Brca1-mouse-scrnaseq-Lev.

## Acknowledgements

T.L. was supported by the NCI under Award Numbers T32GM136624 and F31CA281331. I.A. was supported by the UCI Medical Scientist Training Program (NIH/NIGMS 1T32GM154637-01), and the NCI under Award Number 5T32CA009054-44 and Award Number 1F30CA306157-01. D.M. was supported by a Canadian Institutes of Health Research Postdoctoral Fellowship award and by a NIH/NCI K99/R00 Pathway to Independence Award (1K99CA267160-01). T.P.M. was supported by the UCI Medical Scientist Training Program (NIH/NIGMS 1T32GM154637-01) and the CIRM Training Grant (CIRM EDUC4-12822). K.K. was supported by the funding from the NIH/NCI (R01CA234496), NIH/NIGMS (R01GM147741) and by an Emerging Leader Award from the Mark Foundation for Cancer Research. This work was supported by an NIH/NCI grants (R01CA237376-01A1 and 1P01CA288662-01A1) awarded to D.A.L. This work was also supported in part by an Opportunity Award (U54CA217378) funded by the NCI to the University of California, Irvine, Center for Complex Biological Systems (CCBS; NCI, grant no. U54CA217378) to D.M., T.L., K.K. and D.A.L. This work utilized the infrastructure for high-performance and high-throughput computing, research data storage and analysis, and scientific software tool integration built, operated, and updated by the Research Cyberinfrastructure Center (RCIC) at the University of California, Irvine (UCI). The RCIC provides cluster-based systems, application software, and scalable storage to directly support the UCI research community (https://rcic.uci.edu). We would like to acknowledge Pauline Nguyen and Vanessa Scarfone for all their help in the UCI Flow Cytometry Core. Additional thanks to Melanie Oakes, Quy Nguyen, Jie “Jenny” Wu, Ivan Chang and the UCI Genomics High Throughput Facility staff for assistance with the RNA-sequencing. The development of the Biorepository and generation of the data was supported by the Gray Foundation and Basser Center for BRCA. CIMBA is supported by funding from Cancer Research UK programme grant (PPRPGM-Nov20\100002) the National Health and Medical Research Council of Australia (NHMRC) and the Gray Foundation. We also thank Walid Khaled for insightful thoughts and feedback on the conclusions of the study. Schematics were made using Biorender. The results published here are in part based upon data generated by the TCGA Research Network: https://www.cancer.gov/tcga.

## Author contributions

D.A.L. conceptualized the study. The investigation was conducted by T.L., A.L.P.A.S., I.A. and D.M. The methodology was developed by T.L., A.L.P.A.S., S.M., M.S.L., A.L., T.P.M., D.M., X.L., and X.S.C. Data acquisition was performed by T.L., D.P., A.L.P.A.S., I.A., A.N., K.L.N. and K.K. Data curation was performed by T.L., A.L.P.A.S., I.A. and D.P. Formal analysis was carried out by X.S.C. and T.L. The original manuscript was drafted by T.L. and D.A.L., and all authors contributed to the review and editing of the manuscript. Supervision of the project was provided by K.K., J.L., A.D.L. and D.A.L. Project administration was managed by S.M. and J.G.G. Resources were provided by D.A.L., K.L.N., S.M.D., G.C.-T., A.A., H.T., K.O., P.L.M., P.A.J., A.K.G. and funding was acquired by T.L. and D.A.L.

## Competing interests

The authors declare no competing interests.

**Supplementary Figure 1 | Technical validation and quality control of *Wap-Cre* and *Blg-Cre* scRNA-seq datasets.**

**a**, UMAP projection of the *Wap-Cre* dataset. Cells are colored by sample of origin, with each sample annotated by mouse ID, mammary gland number (1–5) and, where applicable, tumor ID and diameter (mm).

**b–d**, Violin plots with overlaid box plots showing the distribution of the number of detected genes (nFeature_RNA; **b**), total transcript counts (nCount_RNA; **c**) and the percentage of mitochondrial reads (**d**) for each *Wap-Cre* sample. Colors correspond to the sample labels in **a**. Center line, median; box limits, upper and lower quartiles; whiskers, 1.5x the interquartile range. **e**, UMAP projection of *Wap-Cre* cells colored by cell-cycle phase (G1, S or G2M) as assigned by the score_genes_cell_cycle function in Scanpy, utilizing the canonical S and G2/M phase marker gene lists defined in Seurat.

**f**, UMAP re-embedding of the *Wap-Cre* sub-analysis (as shown in **Fig. 1h**) colored by sample of origin as defined in **a**.

**g**, Expression of the proliferation marker *Mki67* visualized on the *Wap-Cre* subset UMAP.

**h**, Schematic of the *Blg-Cre Brca1* mouse model genetics.

**i**, UMAP projection of the *Blg-Cre* dataset, with cells colored by individual sample of origin.

**j**, Stacked bar plot showing the concordance between original cell-type annotations and the re-annotated cell types generated using the pipeline for **Fig. 1e**.

**k**, UMAP re-embedding of the *Blg-Cre* sub-analysis (as shown in Fig. 1k) colored by sample of origin from **i**.

**l**, Expression of *Mki67* visualized on the *Blg-Cre* subset UMAP.

**Supplementary Figure 3 | Comprehensive genotyping and single-cell TNBC subtype marker profiles.**

**a**, Single-cell genotyping (MAESTER) results for the *Wap-Cre* and *Blg-Cre* models. UMAP projection (left) and non-normalized stacked bar plots (right) illustrate the distribution of *Trp53* genotypes (proxy for *Brca1* recombination). Bar heights represent raw cell counts across cell types, experimental conditions and ploidy states. Cells for which a genotype could not be determined are indicated in gray.

**b**, Dot plots showing expression of canonical Bmyo, LumSec and derived TNBC-M marker genes (defined in **Fig. 2** and **Methods**). Data are stratified by single-cell TNBC subtype-like status for wild-type (top) and mutant (bottom) samples. Color intensity represents mean expression level; dot size represents the percentage of cells expressing the gene.

**Supplementary Figure 5 | Validation of developmental atlas integration and reference mapping accuracy.**

**a**, UMAP projection of the Dev-atlas reference (*n* = 95,483 cells). Cells colored by developmental stage.

**b**, UMAP projections of the joint latent space generated by scANVI integration, demonstrating mixing and the absence of batch effects.

**c,** Bar graph displaying the percentage of agreement between original (de novo clustered) and Dev-atlas-defined (scANVI co-clustered) cell populations in the *Wap-Cre* dataset. This metric validates the high fidelity of the joint embedding approach.

**Supplementary Figure 6 | Replicate spatial profiling, region-of-interest annotations, and antibody validation for CODEX multiplex immunofluorescence.**

**a,c,e,** Multiplex immunostained overview sections across all independent biological replicates of *Wap-Cre* mammary tissues during disease progression: wild-type (a), mutant (c) and tumor (e). For each replicate, raw immunofluorescence images exported from QuPath (top) are paired with corresponding spatial cell-segmentation maps rendered via Seurat ImageDimPlot (bottom). Pseudocolored images depict key cell-type lineage markers as defined in **Fig. 6b**. Scale bar, 500 μm. Circles indicate designated $1 mm² regions of interest (ROIs) analyzed in **Fig. 6e** and Extended Data **Fig. 6b**.

**b,d,f,** Non-normalized stacked bar plots displaying raw cell counts across experimental conditions for wild-type (**b**), mutant (**d**) and tumor (**f**) samples, stratified by cell-type composition inside versus outside designated ROIs. Bar heights represent total cell counts.

**Supplementary Figure 7 | Validation of metastatic cell enrichment and atlas integration. a**, FACS plots of control samples and stains demonstrating the robustness of assay.

**b**, FACS plots for all samples and replicates used in the experimental metastasis study, demonstrating consistent isolation of the H2-K^b+^ H2-K^d-^ donor cell fraction.

**c,** UMAP projection of the joint latent space, with cells colored by individual sample ID for all primary and metastatic replicates, demonstrating mixing and the absence of batch effects.

**Supplementary Figure 8 | Validation of cross-species reference mapping and marker conservation.**

**a**, UMAP projection of the HBCA reference (*n* = 165,557 cells) colored by patient ID.

**b**, UMAP projections of the joint latent space (HBCA and mouse models) colored by individual sample ID.

**c,d,** Bar graph displaying the percentage of agreement between original (de novo clustered) and HBCA-defined (mapped) cell populations in the HBCA **(c)** and in wild-type mouse samples **(d).** Separate LumSec cell states were merged into a single LumSec cell population for this validation to ensure cross-species comparability.

**e,** Dot plots showing conserved expression of markers across species. Color intensity represents mean expression level; dot size represents the percentage of cells expressing the gene. Mouse genes were converted to human orthologs.

---

**Extended Data Fig. 1.**
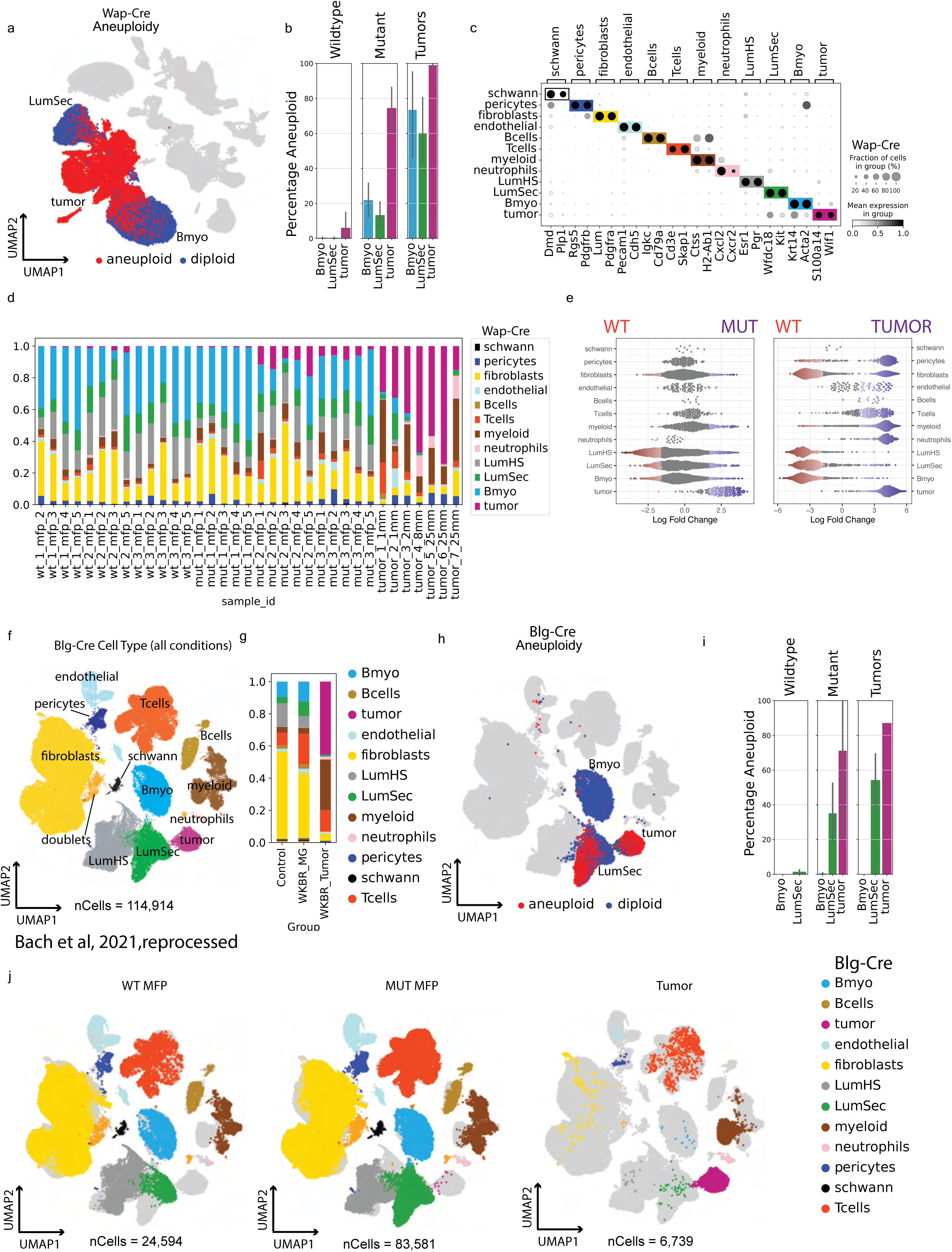
Aneuploidy and cellular composition analysis across *Wap-Cre* and *Blg-Cre* models. **a,** UMAP projection of *Wap-Cre* mammary cells colored by ploidy status as inferred by CopyKAT. **b,** Mean percentage of aneuploid cells within each cell type, stratified by experimental condition (wild-type, mutant and tumor). Error bars represent mean ± s.e.m. for replicates per condition, as defined in Fig. 1c. **c,** Dot plot showing expression of canonical markers across annotated cell types. Dot size represents the percentage of cells expressing the marker; color intensity indicates average expression. **d,** Relative cell-type frequencies for individual *Wap-Cre* samples, including separate mammary glands and tumors. **e,** Beeswarm plot of Milo differential abundance testing, showing shifts in cell-state populations across conditions in the *Wap-Cre* model. **f,g,** UMAP re-embedding **(f)** and relative cell-type frequencies **(g)** for the *Blg-Cre* dataset, processed using the same pipeline as in Fig. 1e**,g**. **h,i,** CopyKAT-inferred ploidy visualized by UMAP **(h)** and mean percentage of aneuploid cells per cell type **(i)** for the *Blg-Cre* model, as described in a,b, with replicates as defined in Fig. 1d. **j,** UMAP projections of the *Blg-Cre* data partitioned by the conditions defined in Fig. 1d.

**Extended Data Fig. 2.**
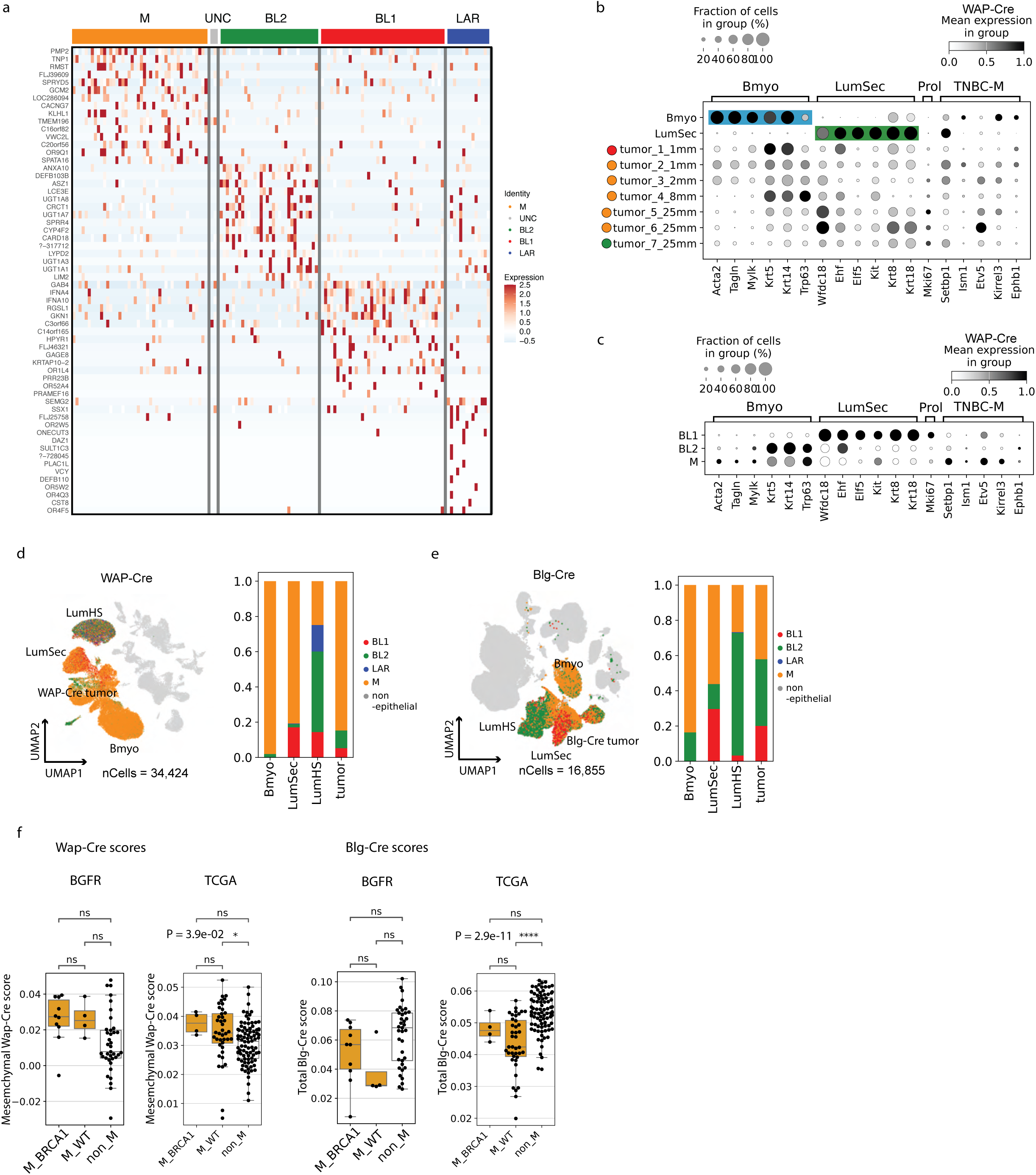
Molecular marker expression and subtype characterization of *Wap-Cre* and *Blg-Cre* models. **a**, Heatmap showing the top 15 differentially expressed genes defining the four human TCGA TNBC subtypes. **b,c**, Dot plots showing expression of canonical basal-myoepithelial (Bmyo), luminal-secretory (LumSec) and derived TNBC-M marker genes (defined in Fig. 2 and **Methods**). Data are shown for individual *Wap-Cre* tumor samples compared to wild-type Bmyo and LumSec cells (**b**) and for aneuploid tumor cells grouped by single-cell TNBC subtype (**c**). Color intensity represents mean expression level; dot size represents the percentage of cells expressing the gene. **d,e**, Single-cell TNBC subtype-like assignment of total epithelial cells in *Wap-Cre* (**d**) and *Blg-Cre* (**e**) models (*n* = 51,279 cells). UMAP projections (left) and stacked bar plots (right) indicate the relative proportions of each subtype. **f,** Box plots of enrichment scores for the *Wap-Cre* mesenchymal tumor signature and the *Blg-Cre* tumor signature in human clinical samples, as in Fig. 2h**,i**, with the mesenchymal (M) group further sub-stratified by *BRCA1* mutation status. The BGFR (*n* = 51) and TCGA (*n* = 132) cohorts are shown separately, as labeled. Center line, median; box limits, upper and lower quartiles; whiskers, 1.5x the interquartile range. *P* values determined by one-way ANOVA followed by Tukey’s post-hoc test, performed separately within each cohort; cohorts were not compared.

**Extended Data Fig. 3.**
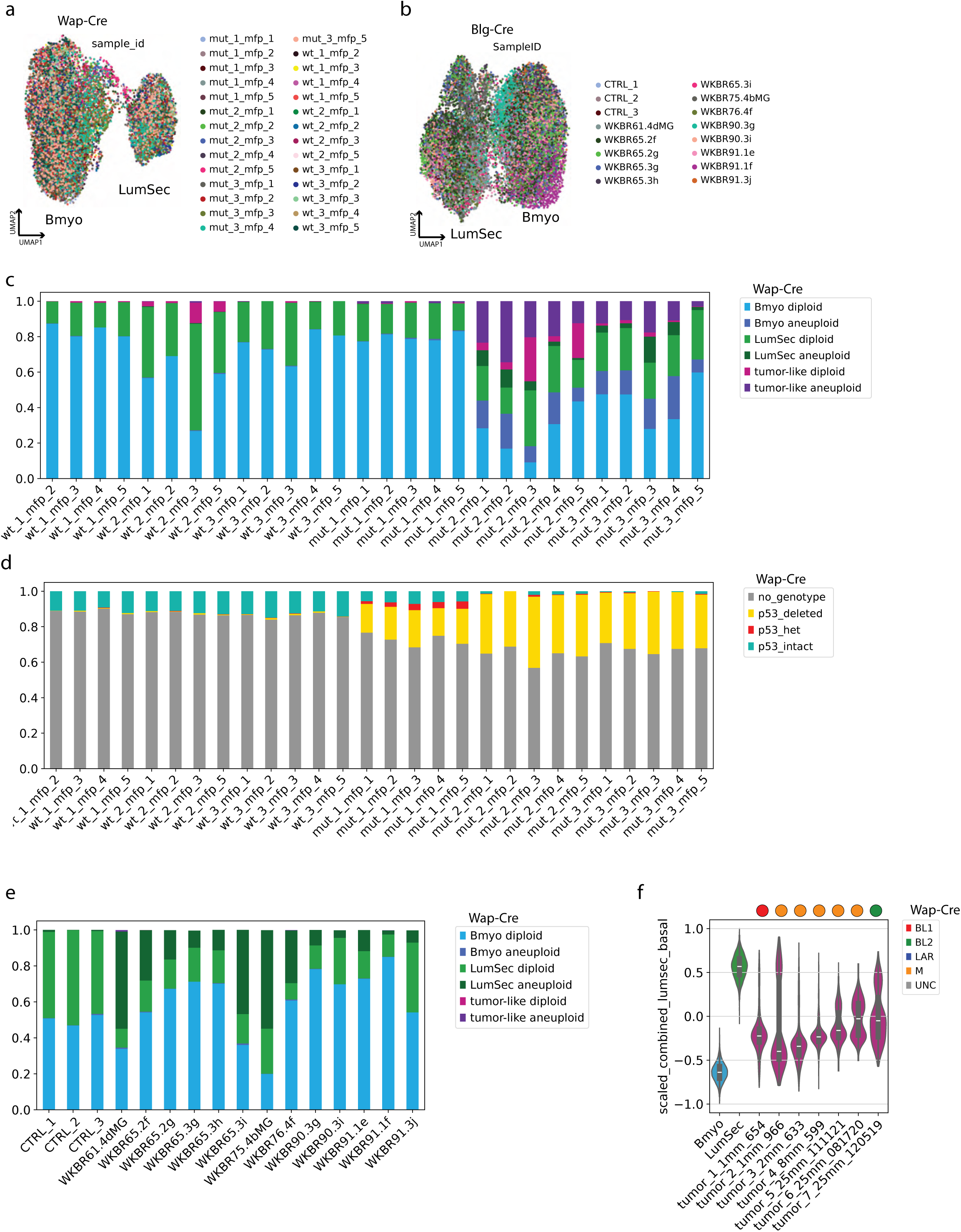
Sample-level distribution of cell types, genotypes and transcriptional scores. **a,b,** UMAP projections of the *Wap-Cre* **(a)** and *Blg-Cre* **(b)** datasets, with cells colored by individual sample of origin. **c,** Relative proportions of epithelial cell types and ploidy status (diploid vs. aneuploid) across *Wap-Cre* samples. **d,** Distribution of single-cell *Trp53* genotypes (MAESTER, used as a proxy for *Brca1* recombination) across *Wap-Cre* samples. Cells without a genotype are indicated in gray. **e,** Relative proportions of epithelial cell types and ploidy status across *Blg-Cre* samples. **f,** Violin plots with overlaid box plots displaying Bmyo–LumSec composite scores for each individual *Wap-Cre* tumor, compared to wild-type control Bmyo and LumSec cells. For box plots, center line represents the median; box limits, upper and lower quartiles; whiskers, 1.5x the interquartile range. Circles above each tumor violin indicate the predicted TNBC subtype based on pseudo-bulk analysis of all cells within that sample, corresponding to the classifications shown in Fig. 2d.

**Extended Data Fig. 4.**
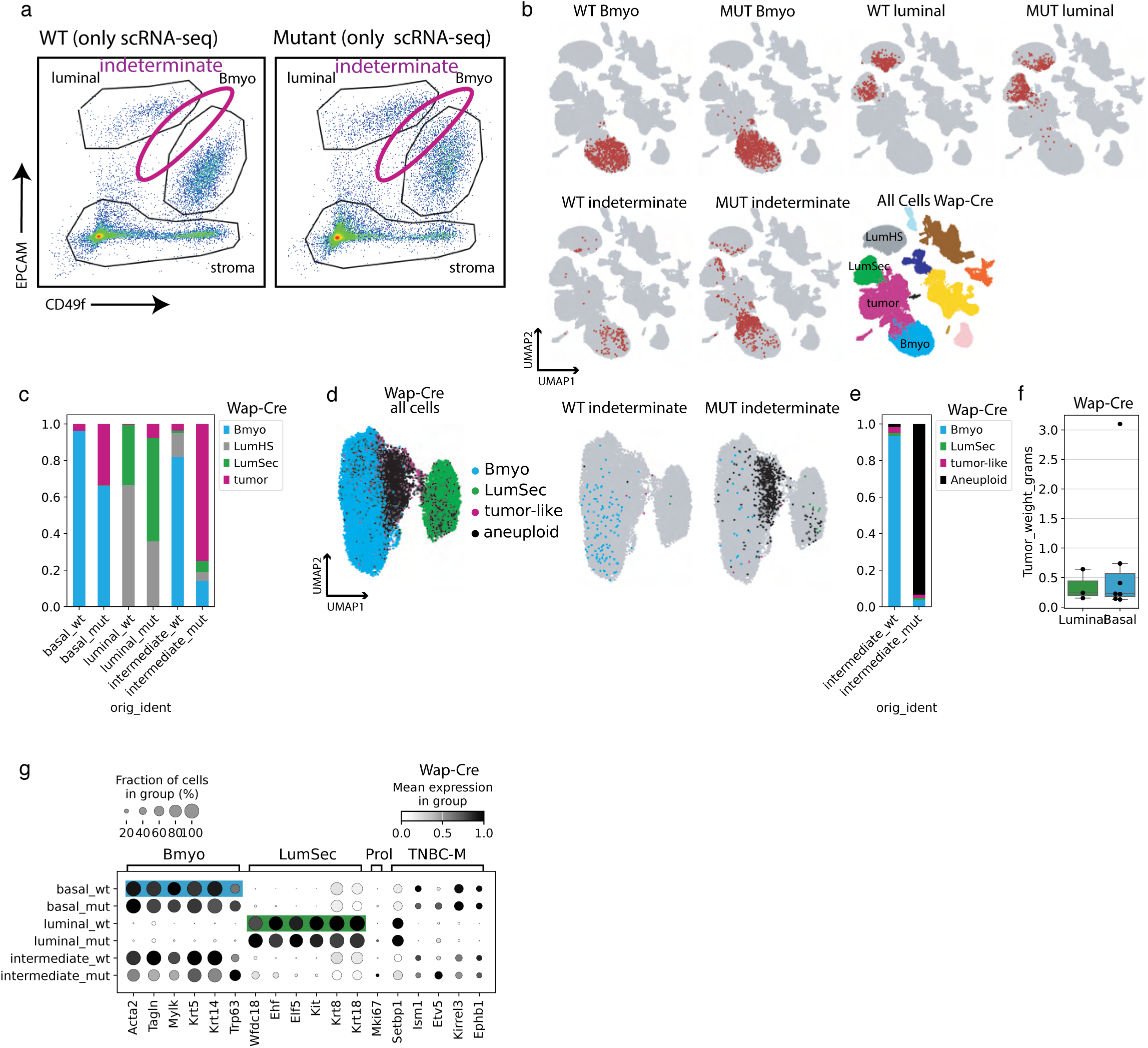
Transcriptomic characterization and tumor-forming potential of indeterminate-lineage mammary epithelial cell fractions. **a,** Representative FACS plots and gating strategy for the isolation of indeterminate-lineage mammary epithelial cell (MEC) subsets from wild-type (left) and mutant (right) donor mice. Indeterminate-lineage cells were defined as Lin^-^ (Cd45^-^) and partitioned using an intermediate gate between canonical basal and luminal populations (Epcam^mid-high^ CD49f^mid-high^). Sorting windows are indicated by magenta ellipses. **b,** Global UMAP projections (as in Fig. 1e) partitioned by genotype and the specific FACS gate of origin, demonstrating the transcriptomic purity of the sorted cell fractions and the inclusion of the LumHS population. **c,** Relative cell-type frequencies across the experimental conditions and gates shown in **a** and Fig. 4b, confirming minimal cross-contamination between basal (Bmyo) and luminal (LumSec and LumHS) identities post-sorting. **d,** UMAP re-embedding of epithelial subsets for all samples (left), wild-type indeterminate (middle) and mutant indeterminate (right) epithelial cells. **e,** Relative cell-type frequencies for the specific conditions and gates shown in the middle and right panels of d. **f,** Box plots showing the mass (grams) of tumors formed in NSG recipients following orthotopic implantation of sorted basal or luminal mutant *Wap-Cre* cell fractions (*n* = 10 tumors; seven basal-derived and three luminal-derived). Center line, median; box limits, upper and lower quartiles; whiskers, 1.5x the interquartile range. **g,** Dot plots showing expression of canonical Bmyo, LumSec and derived TNBC-M marker genes (defined in Fig. 2 and **Methods**). Data are stratified by the experimental conditions and gates shown in **a** and Fig. 4b. Color intensity represents mean expression level; dot size represents the percentage of cells expressing the gene.

**Extended Data Fig. 5.**
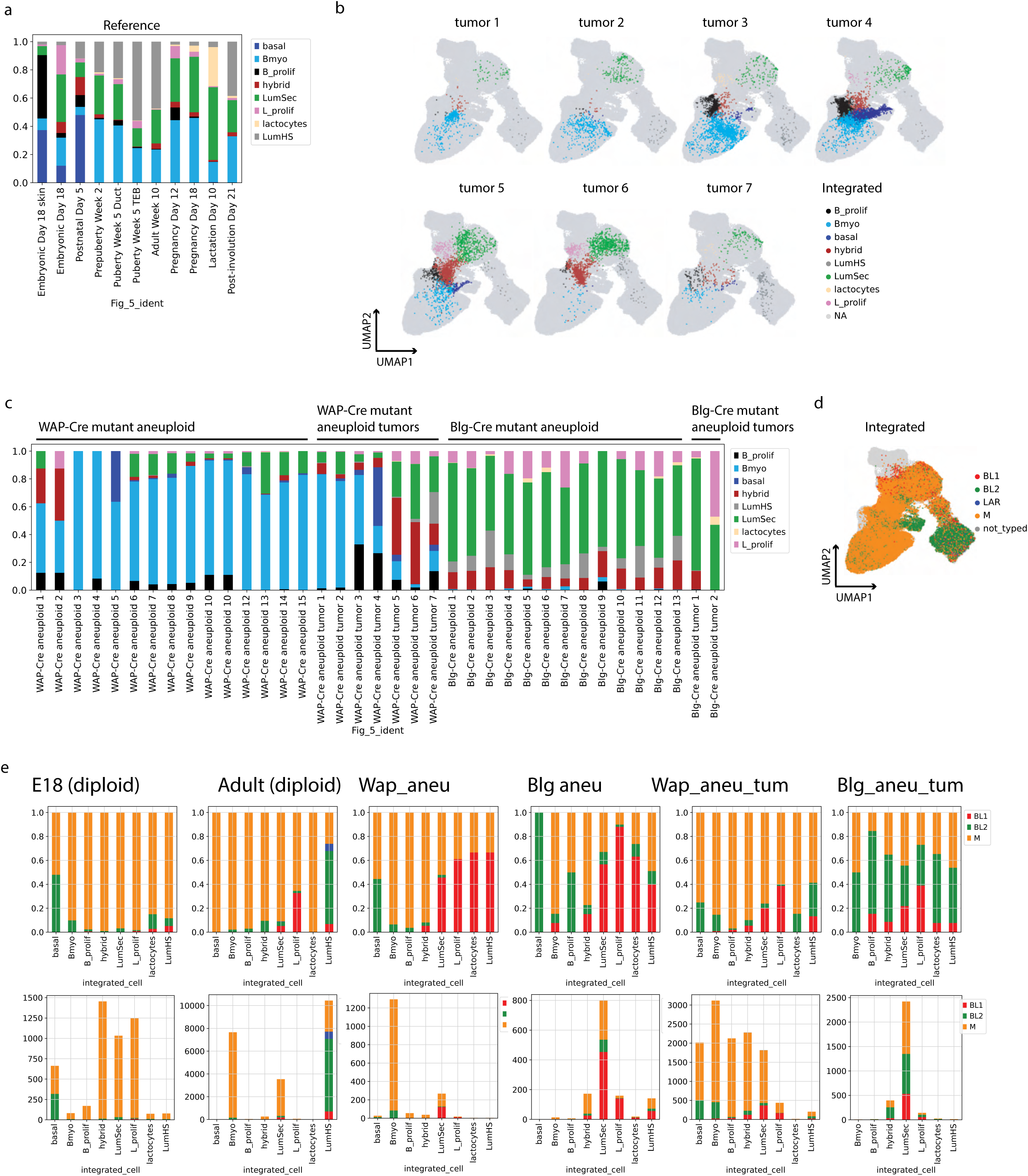
Sample-specific distribution of Dev-atlas-defined cell-states. **a,** Relative cell-type frequencies within the reference Dev-atlas dataset, illustrating the distribution of cell populations identified through re-analysis across the developmental stages in Fig. 5a**,b**. **b,** UMAP projections of the joint latent space generated by scANVI integration of the Dev-atlas and the *Brca1*-deficient datasets. Cells are colored by Dev-atlas-defined cell populations identified through joint scANVI co-clustering and partitioned by individual *Wap-Cre* tumor ID (*n* = 7 tumors). **c,** Relative frequencies of Dev-atlas-defined populations within the aneuploid fraction of individual *Wap-Cre* (*n* = 22) and *Blg-Cre* (*n* = 15) samples. **d,** UMAP projections of the joint latent space, with individual cells colored by assigned human TNBC molecular subtype-like identity. **e,** Distribution of single-cell TNBC subtype-like assignments across Dev-atlas-defined cell populations. Top: Stacked bar plots showing relative frequencies. Bottom: Non-normalized stacked bar plots (bar heights represent raw cell counts).

**Extended Data Fig. 6.**
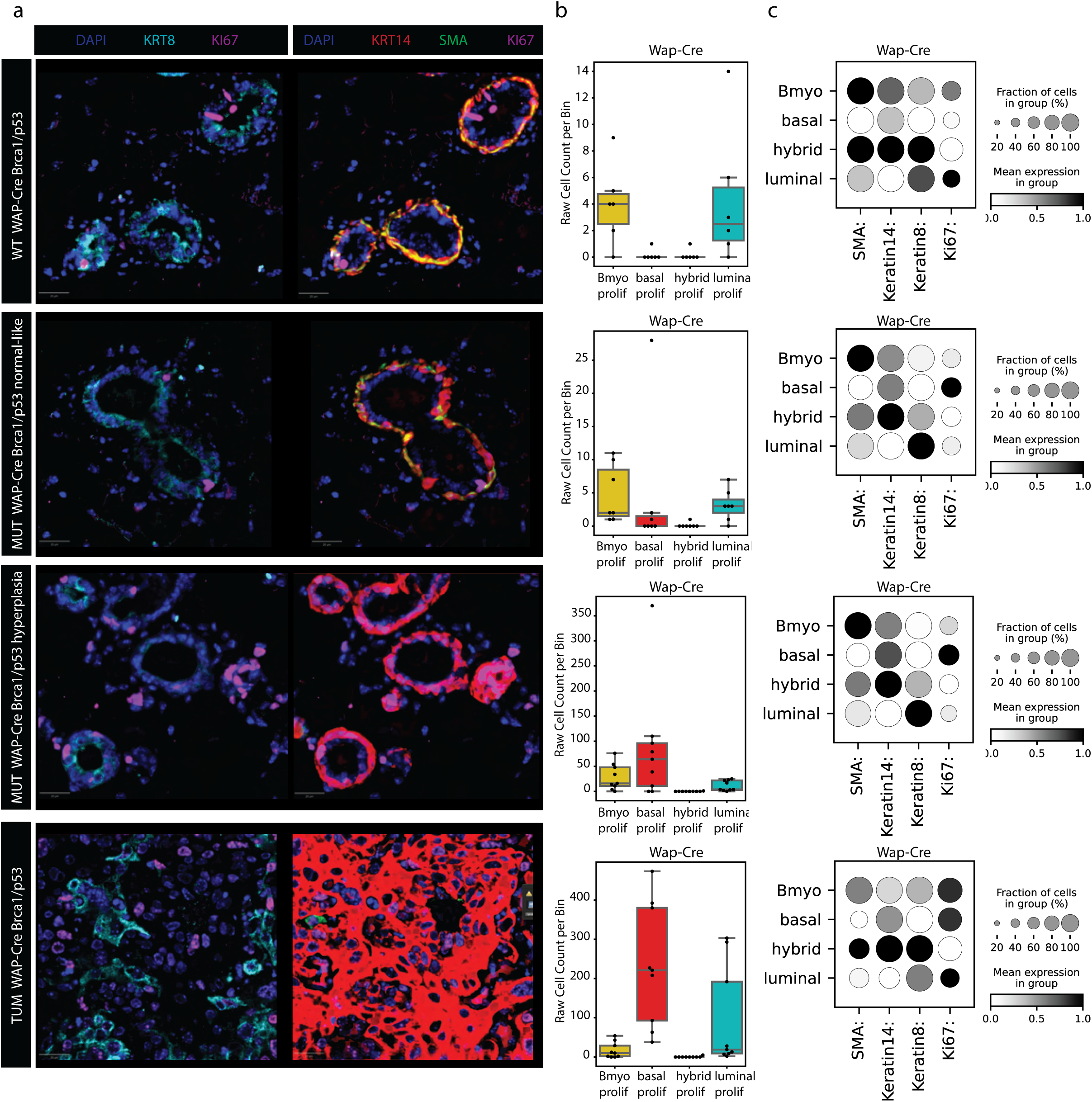
Proliferative dynamics during *Wap-Cre* disease progression. **a,** Representative multiplex immunostained sections of *Wap-Cre* mammary tissues across disease progression: wild-type (top), pre-tumor mutant normal-like (second row), mutant hyperplastic (third row) and tumor (bottom). Pseudocolored images depict key cell-type lineage markers as defined in Fig. 6b, with the addition of the proliferation marker MKI67 (KI67). Scale bar, 20 μm. Data are representative of n = 3 mice per condition, with similar results obtained across n = 3 independent experiments. **b,** Box plots with overlaid data points showing the density (cells per ROI) of proliferating (MKI67⁺) segmented cell types across the experimental conditions described in **a**. Regions of interest (ROIs; 1 mm²) for normal-like and hyperplastic tissues were defined as described in Methods (all ROIs shown in **Supplementary** Fig. 6). Center line, median; box limits, upper and lower quartiles; whiskers, 1.5x the interquartile range. **c,** Dot plot showing protein expression of canonical lineage and proliferation markers across the experimental conditions described in **a**. Color intensity represents mean expression level; dot size represents the percentage of cells expressing the protein.

**Extended Data Fig. 7.**
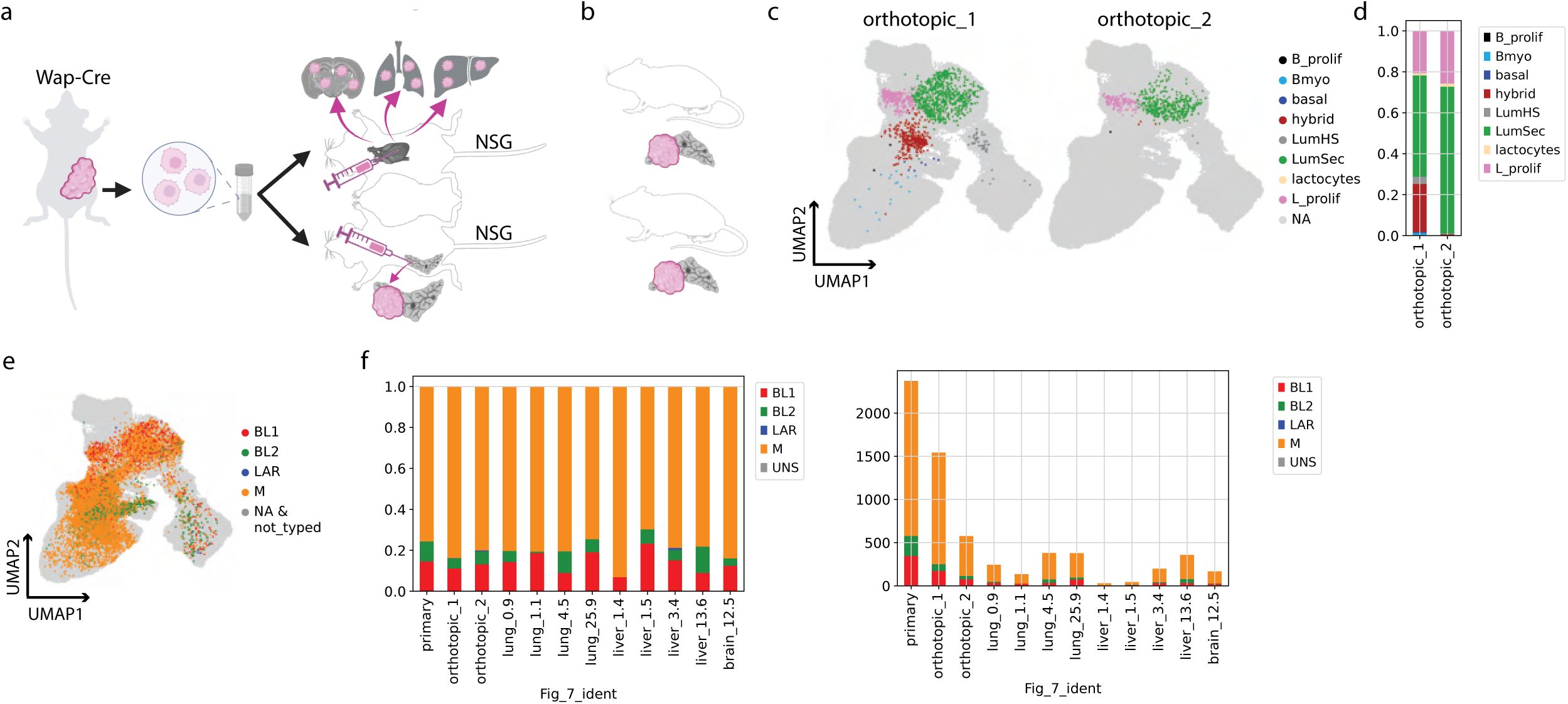
Orthotopic transplantation and TNBC subtyping of metastatic *Wap-Cre Brca1*-deficient cells. **a,** Experimental workflow for orthotopic tumor transplantation. Dissociated primary tumor cells from *Wap-Cre* donors were injected into the mammary fat pads of female NSG recipient mice (*n* = 2 mice; 250,000 cells per injection). **b,** Schematic representation of resulting orthotopic tumors. **c,** UMAP projections of the joint latent space for the orthotopic tumors (n = 2,119 cells), colored by Dev-atlas-defined cell populations identified through joint scANVI co-clustering. **d,** Relative frequencies of Dev-atlas-defined cell populations across the orthotopic samples shown in c. **e,** UMAP projection of the joint latent space for all samples in the metastasis study (n = 6,422 cells), with cells colored by computationally assigned human TNBC molecular subtype. **f,** Distribution of single-cell TNBC subtypes across the metastasis experiment. Left: Stacked bar plots showing relative frequencies. Right: Non-normalized stacked bar plots (bar heights represent raw cell counts).

**Extended Data Fig. 8.**
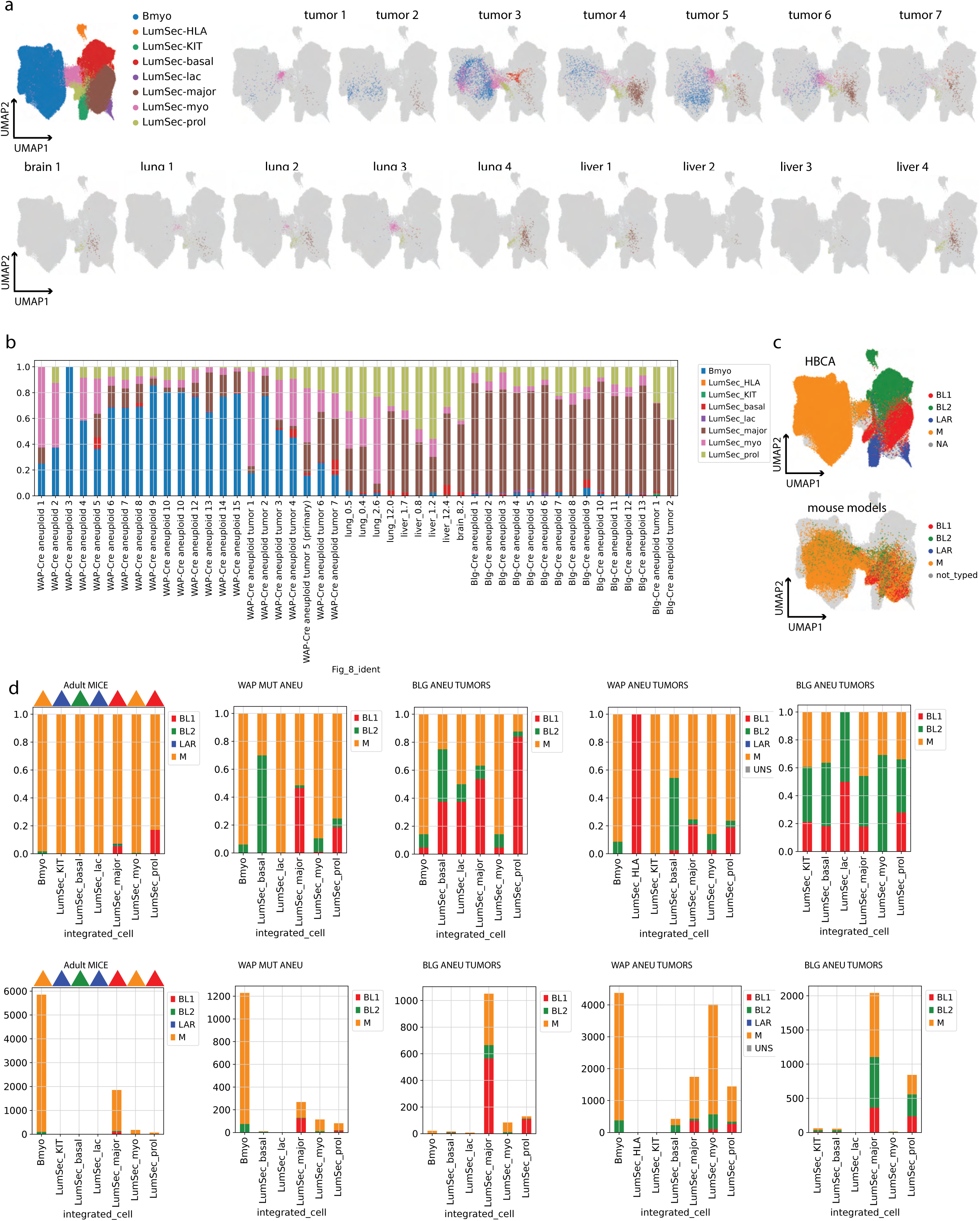
Characterization of tumor heterogeneity and TNBC subtyping in the cross-species integrated atlas. **a,** UMAP projections of the joint latent space (HBCA and mouse models). Cells are colored by HBCA-defined cell populations and partitioned by tumor and metastatic samples. **b,** Relative cell-type frequencies illustrating the distribution of HBCA-defined populations within *Brca1*-deficient samples. **c,** UMAP projections of the joint latent space, colored by assigned human TNBC molecular subtype-like identities. Top: HBCA cells colored by predicted TNBC subtype-like identity based on pseudo-bulk analysis of cell states. Bottom: Mouse cells colored by single-cell TNBC subtype-like identity. **d,** Distribution of single-cell TNBC subtype-like assignments across HBCA-defined cell populations. Top: Stacked bar plots showing relative frequencies. Bottom: Non-normalized stacked bar plots (bar heights represent raw cell counts). Triangles above the “adult mouse” bars indicate the predicted TNBC subtype derived from human cell state pseudo-bulks depicted in **c**.

## References

1. Kuchenbaecker, K. B. et al. Risks of Breast, Ovarian, and Contralateral Breast Cancer for BRCA1 and BRCA2 Mutation Carriers. JAMA 317, 2402–2416 (2017).

2. Moynahan, M. E., Chiu, J. W., Koller, B. H. & Jasin, M. Brca1 controls homology-directed DNA repair. Mol. Cell 4, 511–518 (1999).

3. Cortez, D., Wang, Y., Qin, J. & Elledge, S. J. Requirement of ATM-dependent phosphorylation of brca1 in the DNA damage response to double-strand breaks. Science 286, 1162–1166 (1999).

4. Xu, X. et al. Centrosome amplification and a defective G2-M cell cycle checkpoint induce genetic instability in BRCA1 exon 11 isoform-deficient cells. Mol. Cell 3, 389–395 (1999).

5. Ceccaldi, R. et al. Homologous-recombination-deficient tumours are dependent on Polθ-mediated repair. Nature 518, 258–262 (2015).

6. Nik-Zainal, S. et al. Landscape of somatic mutations in 560 breast cancer whole-genome sequences. Nature 534, 47–54 (2016).

7. Davies, H. et al. HRDetect is a predictor of BRCA1 and BRCA2 deficiency based on mutational signatures. Nat. Med. 23, 517–525 (2017).

8. Popova, T. et al. Ploidy and large-scale genomic instability consistently identify basal-like breast carcinomas with BRCA1/2 inactivation. Cancer Res. 72, 5454–5462 (2012).

9. Calvo, V. & Beato, M. BRCA1 counteracts progesterone action by ubiquitination leading to progesterone receptor degradation and epigenetic silencing of target promoters. Cancer Res. 71, 3422–3431 (2011).

10. Dizin, E. & Irminger-Finger, I. Negative feedback loop of BRCA1-BARD1 ubiquitin ligase on estrogen receptor alpha stability and activity antagonized by cancer-associated isoform of BARD1. Int. J. Biochem. Cell Biol. 42, 693–700 (2010).

11. Eakin, C. M., MacCoss, M. J., Finney, G. L. & Klevit, R. E. Estrogen receptor α is a putative substrate for the BRCA1 ubiquitin ligase. Proc. Natl. Acad. Sci. U. S. A. 104, 5794–5799 (2007).

12. Ellmann, S. et al. Estrogen and progesterone receptors: from molecular structures to clinical targets. Cell. Mol. Life Sci. CMLS 66, 2405–2426 (2009).

13. Faulds, M. H., Olsen, H., Helguero, L. A., Gustafsson, J.-Å. & Haldosén, L.-A. Estrogen Receptor Functional Activity Changes during Differentiation of Mammary Epithelial Cells. Mol. Endocrinol. 18, 412–421 (2004).

14. Humphreys, R. C., Lydon, J., O’Malley, B. W. & Rosen, J. M. Mammary gland development is mediated by both stromal and epithelial progesterone receptors. Mol. Endocrinol. 11, 801–811 (1997).

15. Feng, Y., Manka, D., Wagner, K.-U. & Khan, S. A. Estrogen receptor-α expression in the mammary epithelium is required for ductal and alveolar morphogenesis in mice. Proc. Natl. Acad. Sci. 104, 14718–14723 (2007).

16. Dent, R. et al. Triple-Negative Breast Cancer: Clinical Features and Patterns of Recurrence. Clin. Cancer Res. 13, 4429–4434 (2007).

17. Bauer, K. R., Brown, M., Cress, R. D., Parise, C. A. & Caggiano, V. Descriptive analysis of estrogen receptor (ER)-negative, progesterone receptor (PR)-negative, and HER2-negative invasive breast cancer, the so-called triple-negative phenotype: a population-based study from the California cancer Registry. Cancer 109, 1721–1728 (2007).

18. Liedtke, C. et al. Response to neoadjuvant therapy and long-term survival in patients with triple-negative breast cancer. J. Clin. Oncol. Off. J. Am. Soc. Clin. Oncol. 26, 1275–1281 (2008).

19. Robson, M. et al. Olaparib for Metastatic Breast Cancer in Patients with a Germline BRCA Mutation. N. Engl. J. Med. 377, 523–533 (2017).

20. Litton, J. K. et al. Talazoparib in Patients with Advanced Breast Cancer and a Germline BRCA Mutation. N. Engl. J. Med. 379, 753–763 (2018).

21. Schmid, P. et al. Pembrolizumab for Early Triple-Negative Breast Cancer. N. Engl. J. Med. 382, 810–821 (2020).

22. Zhou, Z. & Zhou, Q. Immunotherapy resistance in triple-negative breast cancer: Molecular mechanisms, tumor microenvironment, and therapeutic implications. Front. Oncol. 15, 1630464 (2025).

23. Mehta, A. K. et al. Targeting immunosuppressive macrophages overcomes PARP inhibitor resistance in BRCA1-associated triple-negative breast cancer. *Nat*. Cancer 2, 66–82 (2021).

24. Shah, J. B. et al. Analysis of matched primary and recurrent BRCA1/2 mutation-associated tumors identifies recurrence-specific drivers. Nat. Commun. 13, 6728 (2022).

25. Brown, T. J. et al. The identification of reversion mutations in patients with advanced pancreatic cancer and germline or somatic BRCA or PALB2 variants who were treated with maintenance rucaparib. J. Clin. Oncol. 41, 734–734 (2023).

26. Lehmann, B. D. et al. Identification of human triple-negative breast cancer subtypes and preclinical models for selection of targeted therapies. J. Clin. Invest. 121, 2750–2767 (2011).

27. Lehmann, B. D. et al. Refinement of Triple-Negative Breast Cancer Molecular Subtypes: Implications for Neoadjuvant Chemotherapy Selection. PLOS ONE 11, e0157368 (2016).

28. Lehmann, B. D. et al. Multi-omics analysis identifies therapeutic vulnerabilities in triple-negative breast cancer subtypes. Nat. Commun. 12, 6276 (2021).

29. Santonja, A. et al. Triple negative breast cancer subtypes and pathologic complete response rate to neoadjuvant chemotherapy. Oncotarget 9, 26406–26416 (2018).

30. Echavarria, I. et al. Pathological Response in a Triple-Negative Breast Cancer Cohort Treated with Neoadjuvant Carboplatin and Docetaxel According to Lehmann’s Refined Classification. Clin. Cancer Res. 24, 1845–1852 (2018).

31. Masuda, H. et al. Differential Response to Neoadjuvant Chemotherapy Among 7 Triple-Negative Breast Cancer Molecular Subtypes. Clin. Cancer Res. 19, 5533–5540 (2013).

32. Zhu, X. et al. Stability and variability of molecular subtypes: comparative analysis of primary and metastatic triple-negative breast cancer. Cancer Biol. Med. 21, 784–798 (2024).

33. McCarthy, A. et al. A mouse model of basal-like breast carcinoma with metaplastic elements. J. Pathol. 211, 389–398 (2007).

34. Molyneux, G., et al. *BRCA1* Basal-like Breast Cancers Originate from Luminal Epithelial Progenitors and Not from Basal Stem Cells. Cell Stem Cell 7, 403–417 (2010).

35. Bach, K. et al. Time-resolved single-cell analysis of Brca1 associated mammary tumourigenesis reveals aberrant differentiation of luminal progenitors. Nat. Commun. 12, 1502 (2021).

36. Landragin, C. et al. Epigenomic disorder and partial EMT impair luminal progenitor integrity in Brca1-associated breast tumorigenesis. Mol. Cancer 24, 127 (2025).

37. Poole, A. J. et al. Prevention of Brca1-Mediated Mammary Tumorigenesis in Mice by a Progesterone Antagonist. Science 314, 1467–1470 (2006).

38. Dann, E., Henderson, N. C., Teichmann, S. A., Morgan, M. D. & Marioni, J. C. Differential abundance testing on single-cell data using k-nearest neighbor graphs. Nat. Biotechnol. 40, 245–253 (2022).

39. Gray, G. K. et al. Defining breast epithelial cell types in the single-cell era. Dev. Cell 60, 2218–2236 (2025).

40. Reed, A. D. et al. A single-cell atlas enables mapping of homeostatic cellular shifts in the adult human breast. Nat. Genet. 56, 652–662 (2024).

41. Gao, R. et al. Delineating copy number and clonal substructure in human tumors from single-cell transcriptomes. Nat. Biotechnol. 39, 599–608 (2021).

42. Maxwell, K. N. et al. BRCA locus-specific loss of heterozygosity in germline BRCA1 and BRCA2 carriers. Nat. Commun. 8, 319 (2017).

43. Chen, X. et al. TNBCtype: A Subtyping Tool for Triple-Negative Breast Cancer. Cancer Inform. 11, CIN.S9983 (2012).

44. Miller, T. E. et al. Mitochondrial variant enrichment from high-throughput single-cell RNA-seq resolves clonal populations. Nat. Biotechnol. 40, 1030–1034 (2022).

45. Pal, B. et al. Single cell transcriptome atlas of mouse mammary epithelial cells across development. Breast Cancer Res. BCR 23, 69 (2021).

46. Spike, B. T. et al. A Mammary Stem Cell Population Identified and Characterized in Late Embryogenesis Reveals Similarities to Human Breast Cancer. Cell Stem Cell 10, 183–197 (2012).

47. Trejo, C. L., Luna, G., Dravis, C., Spike, B. T. & Wahl, G. M. Lgr5 is a marker for fetal mammary stem cells, but is not essential for stem cell activity or tumorigenesis. Npj Breast Cancer 3, 16 (2017).

48. Giraddi, R. R. et al. Single-Cell Transcriptomes Distinguish Stem Cell State Changes and Lineage Specification Programs in Early Mammary Gland Development. Cell Rep. 24, 1653–1666.e7 (2018).

49. Song, Y. et al. Patterns of recurrence and metastasis in BRCA1/BRCA2-associated breast cancers. Cancer 126, 271–280 (2020).

50. Garber, H. R. et al. Incidence and impact of brain metastasis in patients with hereditary BRCA1 or BRCA2 mutated invasive breast cancer. NPJ Breast Cancer 8, 46 (2022).

51. Kumar, T. et al. A spatially resolved single-cell genomic atlas of the adult human breast. Nature 620, 181–191 (2023).

52. Koboldt, D. C. et al. Comprehensive molecular portraits of human breast tumours. Nature 490, 61–70 (2012).

53. Akamandisa, M. P. et al. Comparative analysis of distinct genomic landscapes in young-onset gBRCA1/2 breast cancer. JCI Insight 11, e203005.

54. Liu, J. et al. Characterization of BRCA1-deficient premalignant tissues and cancers identifies Plekha5 as a tumor metastasis suppressor. Nat. Commun. 11, 4875 (2020).

55. Deome, K. B., Faulkin, L. J., Bern, H. A. & Blair, P. B. Development of mammary tumors from hyperplastic alveolar nodules transplanted into gland-free mammary fat pads of female C3H mice. Cancer Res. 19, 515–520 (1959).

56. 10x Genomics. Cell Ranger Single-Cell Software Suite. 10x Genomics (2022).

57. Wolf, F. A., Angerer, P. & Theis, F. J. SCANPY: large-scale single-cell gene expression data analysis. Genome Biol. 19, 15 (2018).

58. Hao, Y. et al. Dictionary learning for integrative, multimodal and scalable single-cell analysis. Nat. Biotechnol. 42, 293–304 (2024).

59. Silkwood, K. et al. Leveraging gene correlations in single cell transcriptomic data. BMC Bioinformatics 25, 305 (2024).

60. Dollinger, E. P., Silkwood, K., Atwood, S., Nie, Q. & Lander, A. D. Statistically principled feature selection for single cell transcriptomics. BMC Bioinformatics 26, 238 (2025).

61. McGinnis, C. S., Murrow, L. M. & Gartner, Z. J. DoubletFinder: Doublet Detection in Single-Cell RNA Sequencing Data Using Artificial Nearest Neighbors. Cell Syst. 8, 329–337.e4 (2019).

62. Traag, V. A., Waltman, L. & van Eck, N. J. From Louvain to Leiden: guaranteeing well-connected communities. Sci. Rep. 9, 5233 (2019).

63. McInnes, L., Healy, J., Saul, N. & Großberger, L. UMAP: Uniform Manifold Approximation and Projection. J. Open Source Softw. 3, 861 (2018).

64. Lopez, R., Regier, J., Cole, M. B., Jordan, M. I. & Yosef, N. Deep generative modeling for single-cell transcriptomics. Nat. Methods 15, 1053–1058 (2018).

65. Gayoso, A. et al. A Python library for probabilistic analysis of single-cell omics data. Nat. Biotechnol. 40, 163–166 (2022).

66. Robinson, M. D., McCarthy, D. J. & Smyth, G. K. edgeR: a Bioconductor package for differential expression analysis of digital gene expression data. Bioinformatics 26, 139–140 (2010).

67. Blake, J. A. et al. Mouse Genome Database (MGD): Knowledgebase for mouse–human comparative biology. Nucleic Acids Res. 49, D981–D987 (2021).

68. Virtanen, P. et al. SciPy 1.0: fundamental algorithms for scientific computing in Python. Nat. Methods 17, 261–272 (2020).

69. Vallat, R. Pingouin: statistics in Python. J. Open Source Softw. 3, 1026 (2018).

70. Xu, C. et al. Probabilistic harmonization and annotation of single-cell transcriptomics data with deep generative models. Mol. Syst. Biol. 17, e9620 (2021).

71. Schmidt, U., Weigert, M., Broaddus, C. & Myers, G. Cell Detection with Star-Convex Polygons. in Medical Image Computing and Computer Assisted Intervention – MICCAI 2018 (eds Frangi, A. F., Schnabel, J. A., Davatzikos, C., Alberola-López, C. & Fichtinger, G.) vol. 11071 265–273 (Springer International Publishing, Cham, 2018).

72. qupath/qupath-extension-stardist. QuPath (2026).

