## Supplementary Figures for "Cell-of-origin shapes tumor subtype in *BRCA1*-associated triple-negative breast cancer"

### Supplementary Figure 1

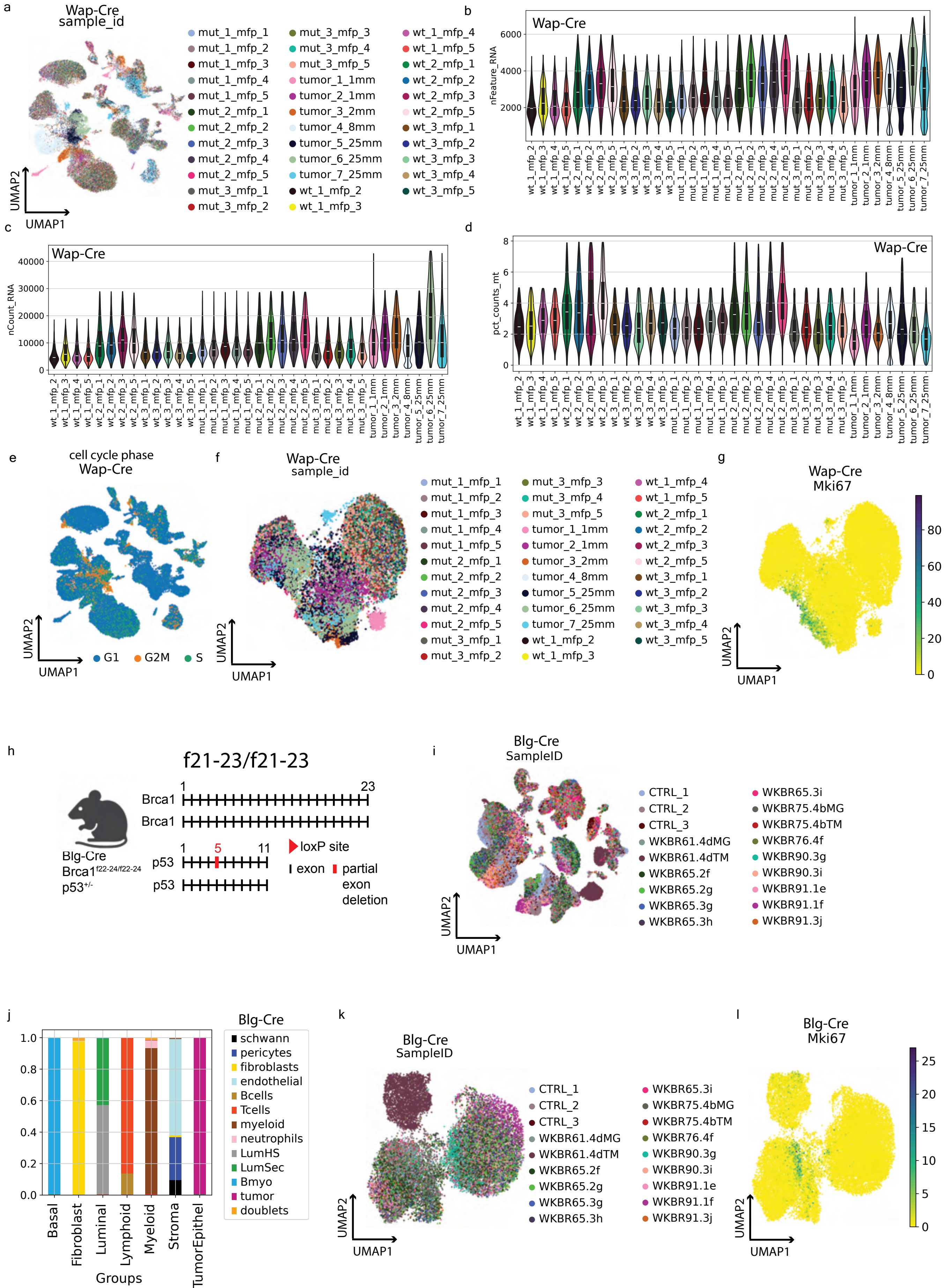

Supplementary Figure 3

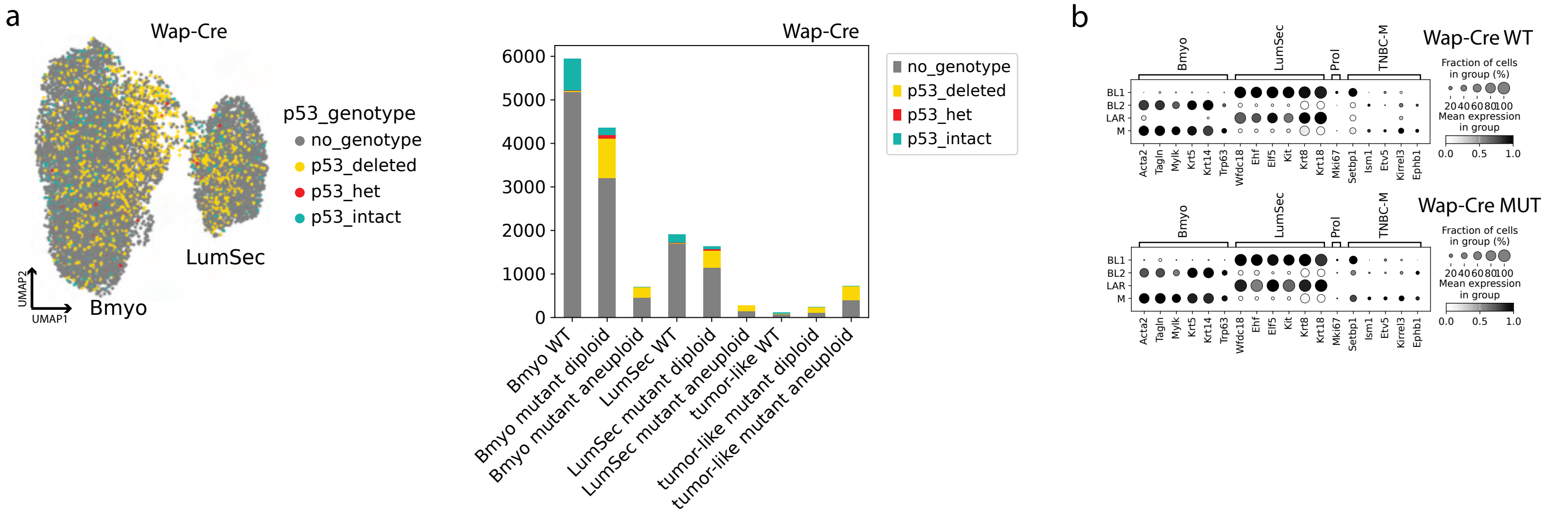

### Supplementary Figure 5

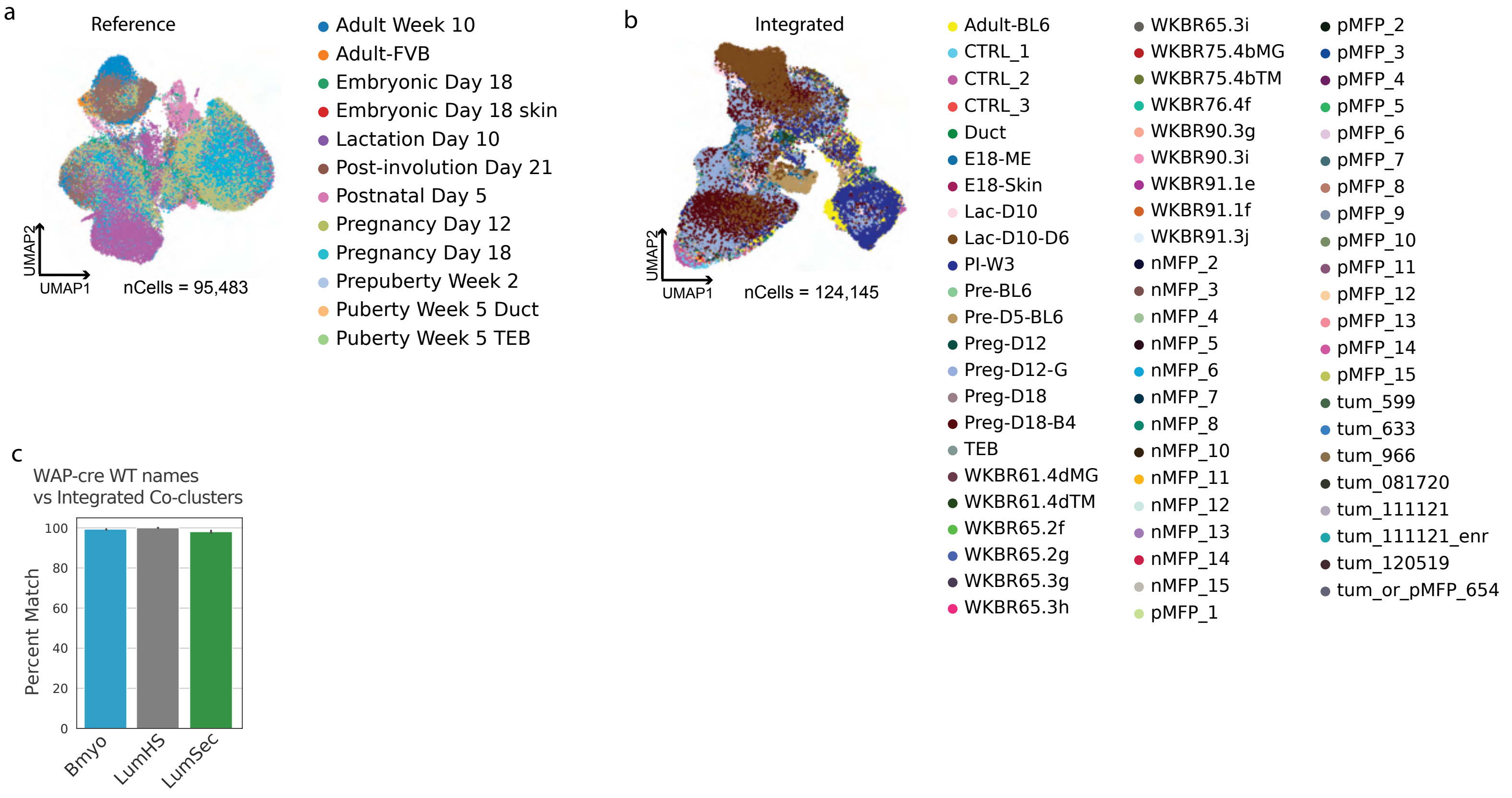

#### Supplementary Figure 6

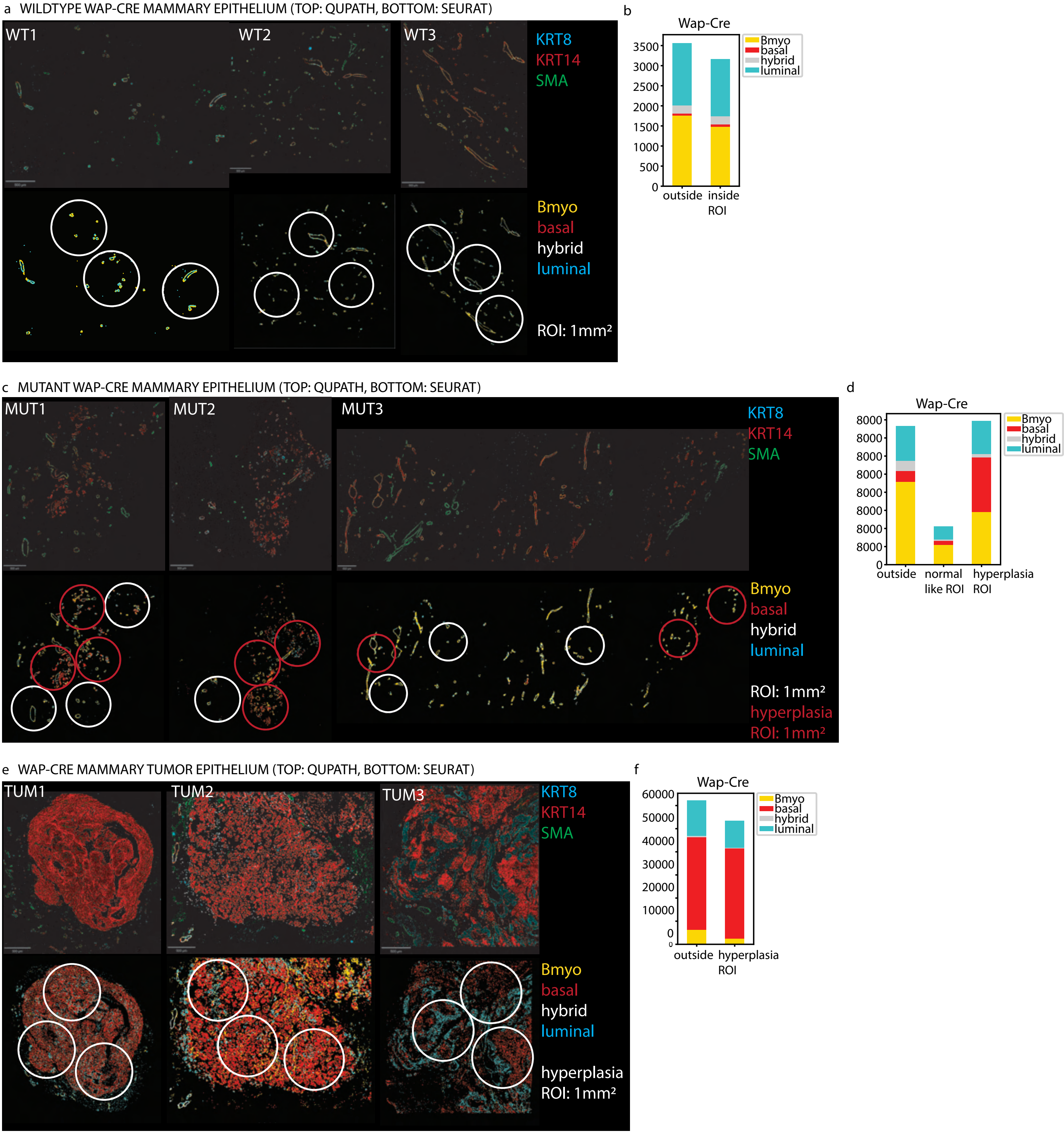

Supplementary Figure 7

a

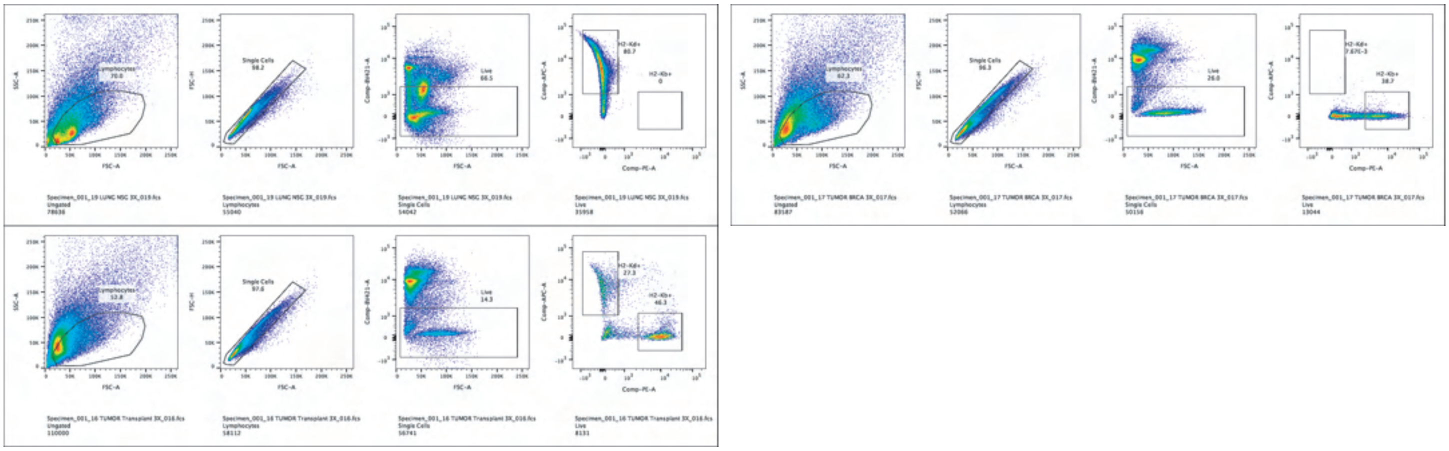

b

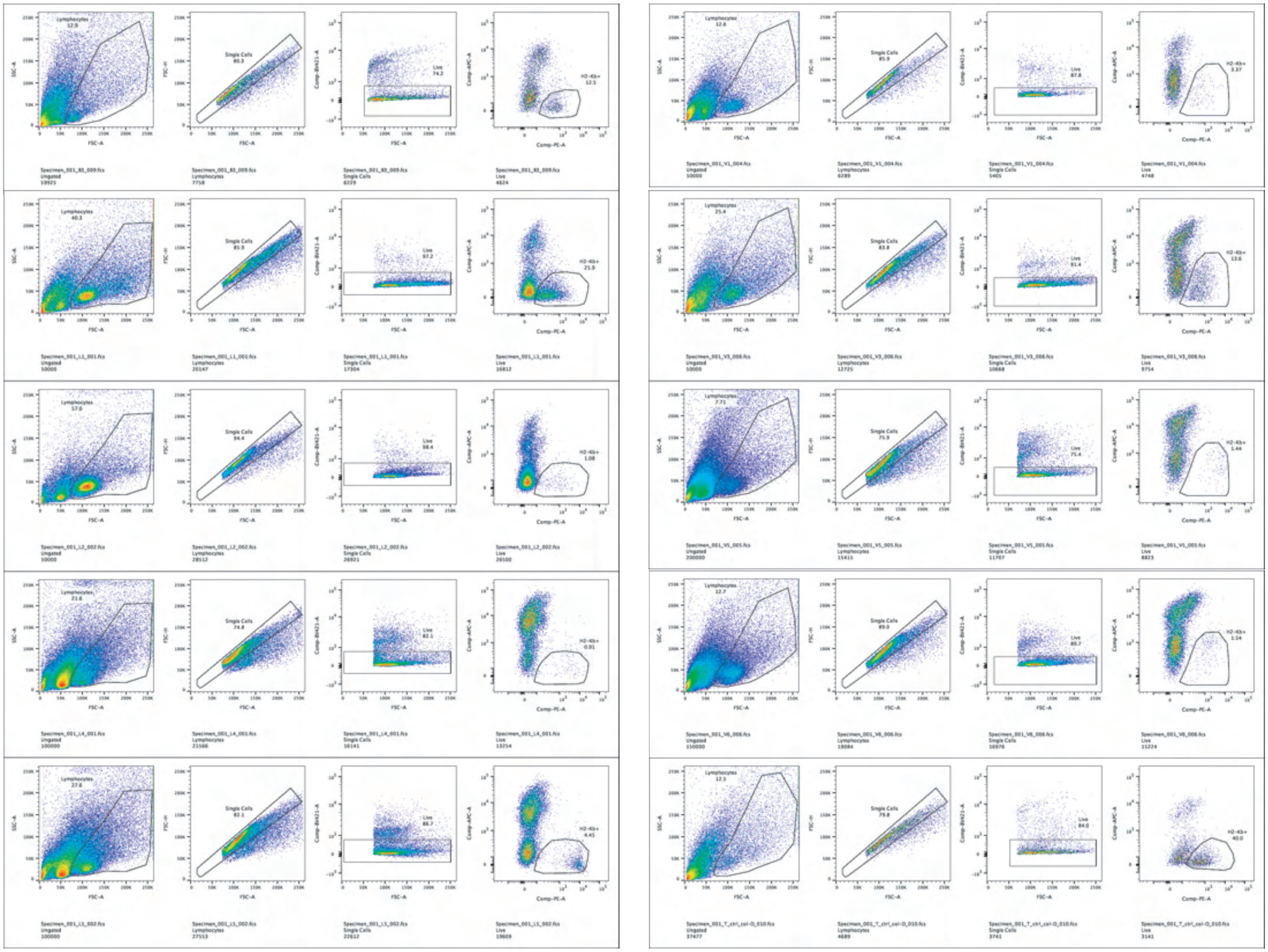

c

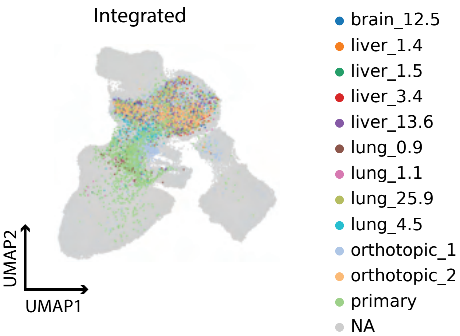

Supplementary Figure 8

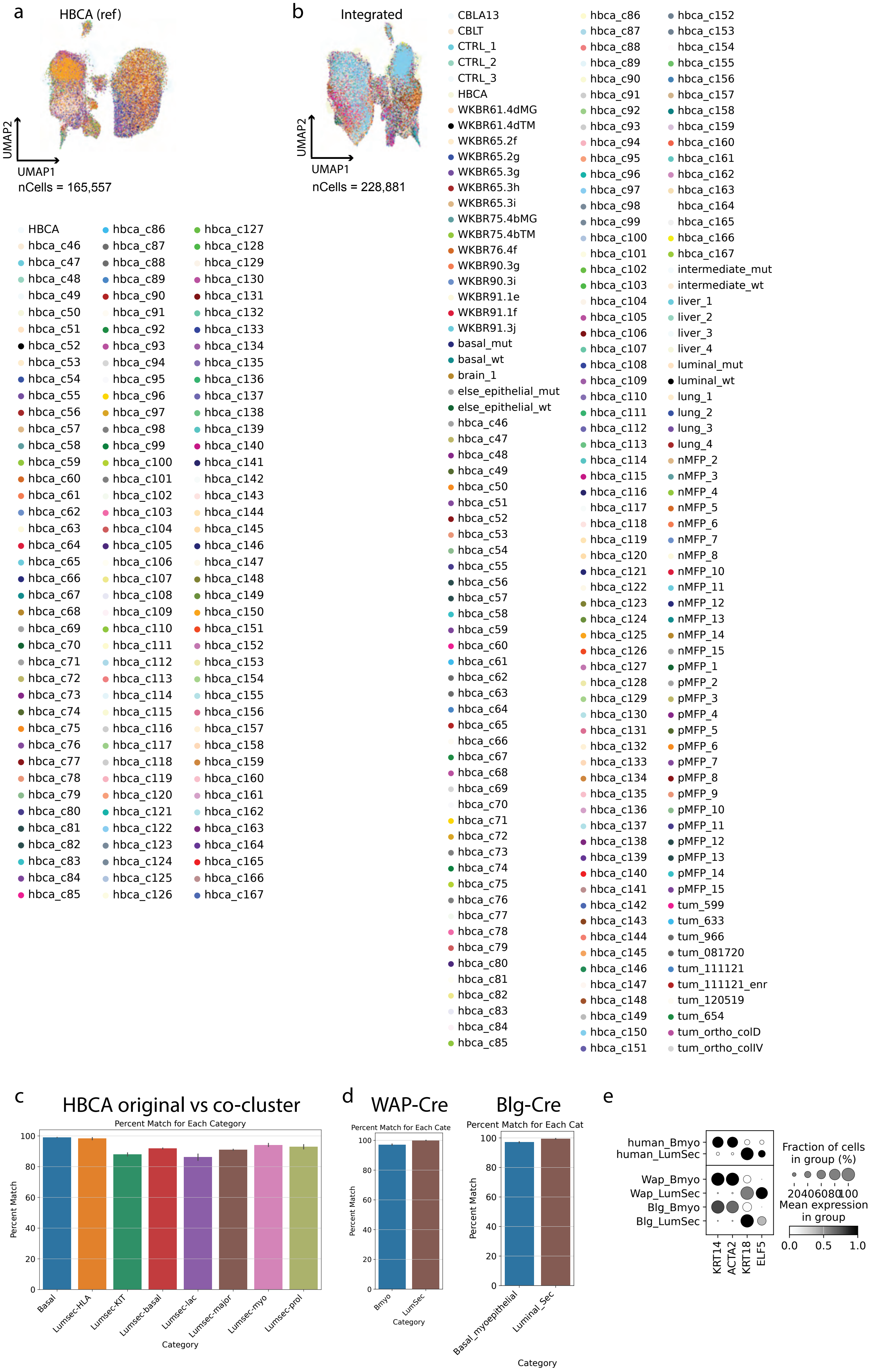
